# Arabidopsis Acyl-CoA Binding Protein 4, ACBP4, functions in developmentally programmed endoreduplication

**DOI:** 10.64898/2026.08.31.748435

**Authors:** Johan Jaenisch, Claudine G. Tahmin, Kai U. Wu, Mary C. Wildermuth

**Affiliations:** Department of Plant & Microbial Biology, University of California, Berkeley, CA 94720-3102

## Abstract

Powdery mildew fungi induce localized endoreduplication, a variant of the cell cycle in which DNA is replicated but cells do not divide, in leaf mesophyll cells underlying the fungal feeding structure. Induced endoreduplication occurs concurrent with powdery mildew (PM) spore production and is associated with enhanced metabolic capacity and flux to lipids. The final ploidy of these cells is highly correlated with fungal spores produced and is the consequence of both basal (developmental) ploidy and PM-induced endoreduplication programs. Herein, we find the Arabidopsis lipid trafficking and regulatory protein ACYL-COA BINDING PROTEIN 4 (ACBP4) enhances PM spore production on Arabidopsis leaves. ACBP4 does not limit plant defense but instead supports basal mesophyll cell ploidy, with decreased final ploidy in cells underlying the fungal feeding structure in *acbp4* mutants compared to wild-type (WT). Leaf epidermal cell size is decreased and stomatal density is increased in *acbp4*, consistent with a role for ACBP4 in developmentally programmed endoreduplication. Moreover, hypocotyl elongation in the dark, which is driven by programmed developmental endoreduplication, shows reduced hypocotyl length, cell length and ploidy in *acbp4* versus WT. Together, our findings establish a novel means by which a plant ACBP promotes cell metabolism and development, with potential applications to agricultural productivity and quality.

## INTRODUCTION

Powdery mildews are obligate biotrophic fungi that infect a wide range of agronomically important monocot and dicot species (Micali et al. 2008; Glawe 2008). Their proliferation is dominated by asexual reproduction, with proliferation occurring every 5-12 days depending on environmental conditions. The life cycle progresses from spore (aka condia) germination on the leaf surface, appressorial formation, penetration of the plant cell wall, formation of a feeding structure (haustorial complex) in the penetrated epidermal cell, surface hyphal network expansion, and asexual reproduction resulting in conidiophores that contain a chain of new spores filled with lipid droplets. Similar to other obligate biotrophs of plants, the genomes of powdery mildew fungi exhibit a number of lost or incomplete metabolic pathways (Spanu et al. 2010; Liang et al. 2018). As such, powdery mildew fungi are dependent on their plant host to meet their nutritional requirements with host susceptibility factors required for their growth and reproduction (Lapin & Van den Ackerveken, 2013; Wildermuth et al. 2017).

In wild-type (WT) *Arabidopsis thaliana* plants infected with the powdery mildew *Golovinomyces orontii MGH1* (Gor), mesophyll cells underlying the fungal feeding structure undergo 2-4 cycles of endoreduplication concurrent with asexual reproduction (5+ days post inoculation (dpi)) (Chandran et al. 2010; 2013). Endoreduplication, a variant of the cell cycle in which cellular DNA is replicated but cell division does not occur, results in a doubling of DNA content with each endocycle round accompanied by nuclear and cell expansion. The Gor-induced local increase in plant cell ploidy (DNA content) is associated with enhanced metabolic capacity including increased expression of glycolytic genes and those involved in a shift in primary metabolism to bypass the pyruvate dehydrogenase complex (PDHc) (Chandran 2010, Wildermuth 2010, Lee et al. 2024). Use of the PDHc bypass then directs flux to lipids which accumulate as triacylglycerols (TAGs) used to support Gor spore production (Lee et al. 2024: Xue et al. 2025). These TAGs, visible as lipid droplets, accumulate in mesophyll cells underlying the fungal feeding structure at 5+ dpi and are made by chloroplast-localized DIACYLGLYCEROL TRANSFERASE 3 (DGAT3), at the expense of thylakoid membranes (Xue et al. 2025). To further explore how localized Arabidopsis lipid metabolism is altered to promote fungal spore production, we assessed a potential role for acyl-CoA binding proteins in powdery mildew growth and asexual reproduction.

Acyl-CoA binding proteins (ACBPs), referred to as ACBDs in animals, are present in eukaryotes and some prokaryotes; they contain a conserved acyl-CoA binding domain (ABD) that binds acyl-CoA lipids with varied lipid preferences (Lai & Chye, 2021; Neess et al. 2015). In plants, ACBPs are known to function in lipid metabolism, membrane biogenesis, development, reproduction, and stress tolerance/response (Lai & Chai, 2021; Hamdam et al., 2021; Wundersitz et al. 2025). Soluble ACBPs are known to direct the intracellular trafficking of fatty acyl-CoAs, protecting their cargo from degradation by acyl-hydrolases and lipases, while membrane-bound ACBPs can tether organelles, facilitating efficient lipid transport. The Arabidopsis genome contains six ACBPs, of which half are membrane-bound and two thirds contain additional protein-protein interaction domains (Figure 1A). The class I member, ACBP6, is a small soluble highly conserved protein that consists largely of the acyl-CoA binding domain and is involved in plant stress and development (Chen et al., 2008; Guo et al., 2019: Hsiao et al., 2014, 2015). Arabidopsis class II members, ACBP1 and ACBP2, are membrane tethered and contain ankyrin repeat protein-protein interaction domains; they have been shown to function in oxidative stress, drought response, and stem cuticle formation (Gao et al. 2010; Du et al. 2013; Xue et al. 2014; Schmidt et al. 2018). The class III ACBP3 exhibits extracellular localization; it includes a transmembrane domain and is proposed to function in cell-to-cell signaling and defense against bacterial and necrotrophic fungal infection (Xiao and Chye 2011; Xia et al. 2012). Arabidopsis ACBP4 and ACBP5 in class IV are soluble cytosolic proteins that contain kelch repeat propeller protein-protein interaction domains (Leung et al. 2005); they impact membrane lipid composition and are involved in development, response to hypoxia and drought (Xiao et al., 2008; Hsiao et al. 2014, 2015; Guo et al. 2024).

**Figure 1:**
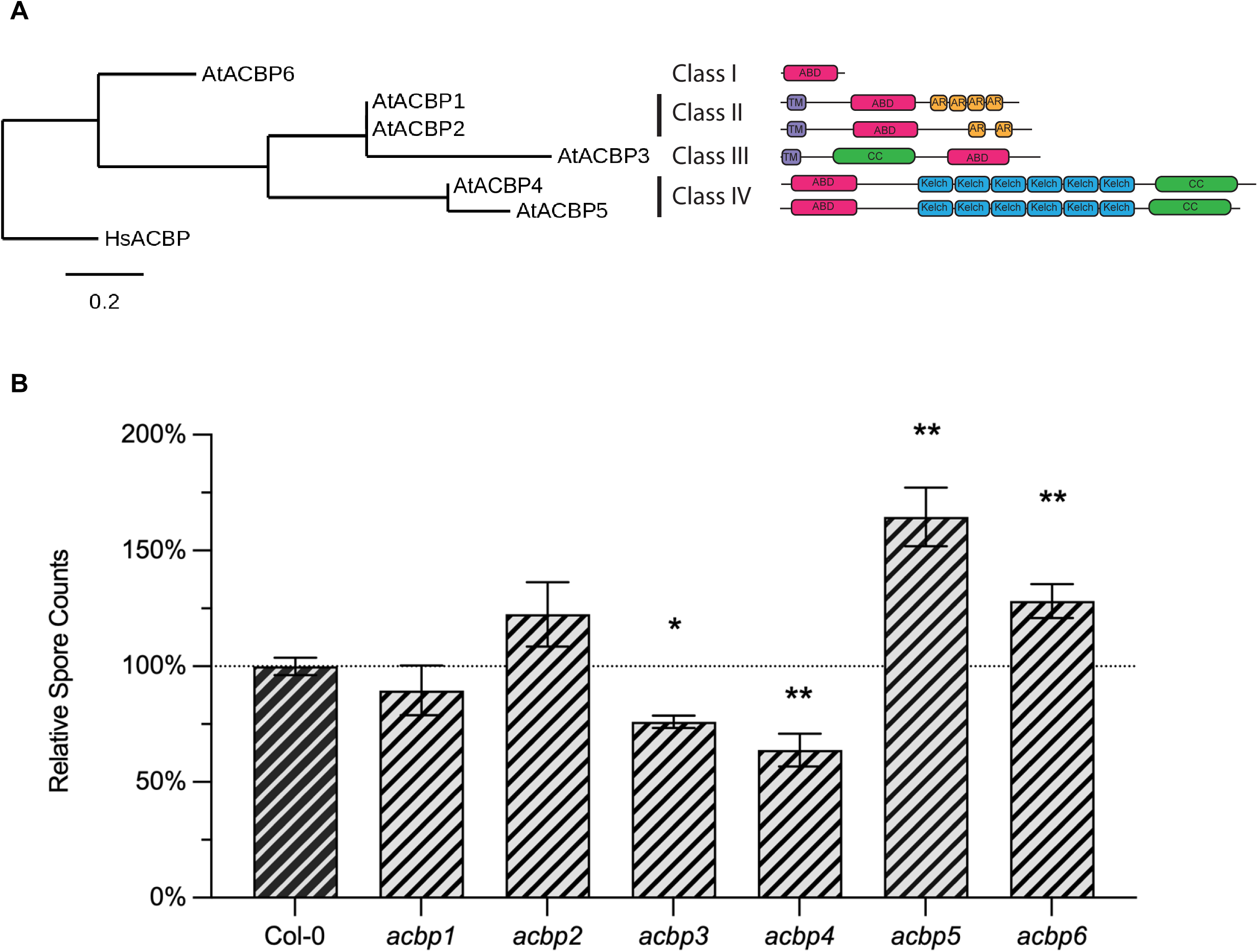
Arabidopsis mutants in acyl-CoA binding protein genes show distinct powdery mildew phenotypes. **A)** Phylogenetic tree of Arabidopsis acyl-CoA binding protein gene family and protein structures. Phylogram using protein sequences of AtACBPs and *Homo sapiens* HsACBD, used as an outgroup, obtained from the Uniprot database, and constructed using Phylogeny.fr (Dereeper et al. 2008). ABD: acyl-CoA binding domain; AR: ankyrin repeats; Kelch: Kelch propeller domain repeats; TM: transmembrane domain; CC: coiled-coil domain. **B)** Powdery mildew reproductive output on AtACBP. Mean Gor spore counts at 9 dpi are normalized to Col-0 wild-type control plants for each genotype at 9 dpi; n ≥ 6. Significant differences from WT were determined using paired 2-tailed t-test (* p ≤ 0.05, ** p ≤ 0.01). Error bars represent ± SEM. Dark gray= Col-0 WT; stripes indicate powdery mildew infection.

In this study, we determine Arabidopsis ACBP4, but not its paralog ACBP5, supports powdery mildew growth and asexual reproduction. This impact is not associated with altered plant defense; instead, it is associated with reduced final ploidy of mesophyll cells underlying the fungal feeding site at 5 dpi in *acbp4* compared to WT. This reduction is due to decreased basal (uninfected) ploidy levels and not compromised Gor-induced endoreduplication. Leaf epidermal cell size and stomatal density are also altered in *acbp4* consistent with a role for ACBP4 in (basal) developmentally programmed endoreduplication. A reduction in hypocotyl elongation in the dark (driven by endoreduplication-dependent cell expansion) for *acbp4* establishes a novel role for a plant ACBP, ACBP4, in facilitating endoreduplication. Understanding novel components of and mechanisms involved in developmentally programmed endoreduplication may allow for increased agricultural productivity and quality for crops including oil and fiber crops, tubers, fleshy fruits and berries that are most likely to benefit from fine-tuning developmentally programmed endoreduplication.

## RESULTS

### Acyl-CoA binding proteins exhibit distinct powdery mildew phenotypes

In order to understand the role of host fatty acid (FA) sensing and trafficking responses during Gor infection, single knockout lines for each of the Arabidopsis ACBP gene family members were screened for powdery mildew proliferation as compared to WT plants (Figure 1A-B). Spore counts at 9 dpi with Gor were used as an endpoint measure of powdery mildew proliferation. Mutants in membrane-bound class II Arabidopsis ACBPs, *acbp1* and *acbp2*, show no significant difference in the amount of spores produced on them relative to WT (Fig. 1B). A mutant in the Class III extracellular-localized and tethered ACBP3 results in a 23% reduction in spore production compared to WT. The mutant in the soluble class I ACBP, ACBP6, exhibits increased spore production (29%) relative to WT. Surprisingly, divergent phenotypes are observed for Class IV mutants *acbp4* and *acbp5*; *acbp4* supports less spore production (39% reduction) while spore production on *acbp5* increases by 47% compared to WT.

### Class IV ACBPs perform distinct roles in the context of powdery mildew infection

Of the Arabidopsis ACBPs, knockouts in the two Class IV ACBPs, ACBP4 and ACBP5, exhibited the greatest alteration in Gor spore production. Previous reports support complementary roles for ACBP4 and ACBP5 in development (reviewed in Du et al., 2016). Therefore, their expression in whole leaves infected with powdery mildew at 5 dpi and in parallel (mock) uninfected (UI) tissue was investigated. Our previously published RNASeq dataset (McRae et al. 2023; analyzed using DESeq2, n=3) shows a minor but statistically significant increase in *ACBP4* expression at 5 dpi versus parallel UI leaves of 1.24x (*adj p-value* ≤ 0.0001) and decrease in *ACBP5* expression of 0.59 (*adj p-value* ≤ 0.05). For confirmation, we performed an independent experiment comparing expression in whole leaves infected with Gor at 5 dpi and UI tissue, with expression analyzed by qPCR. Similar to the RNASeq data, we found *ACBP4* expression is 1.43 higher in infected leaves compared to UI (*adj p-value* ≤ 0.0001), while *ACBP5* expression is reduced (0.73x lower; *adj p-value* ≤ 0.0001) (Fig. 2A). Moreover, eFP-Seq Browser (Sullivan et al. 2019) shows basal expression of *ACBP4* in mature leaves to be ∼10-fold higher than *ACBP5* (log2 normalized kilobase of exon per million mapped reads (RPKM) to average RPKM of controls; Aerial part of 4-week-old plant (SRS360059), Leaf of 4-week-old long-day-grown plant (SRS398009)).

**Figure 2:**
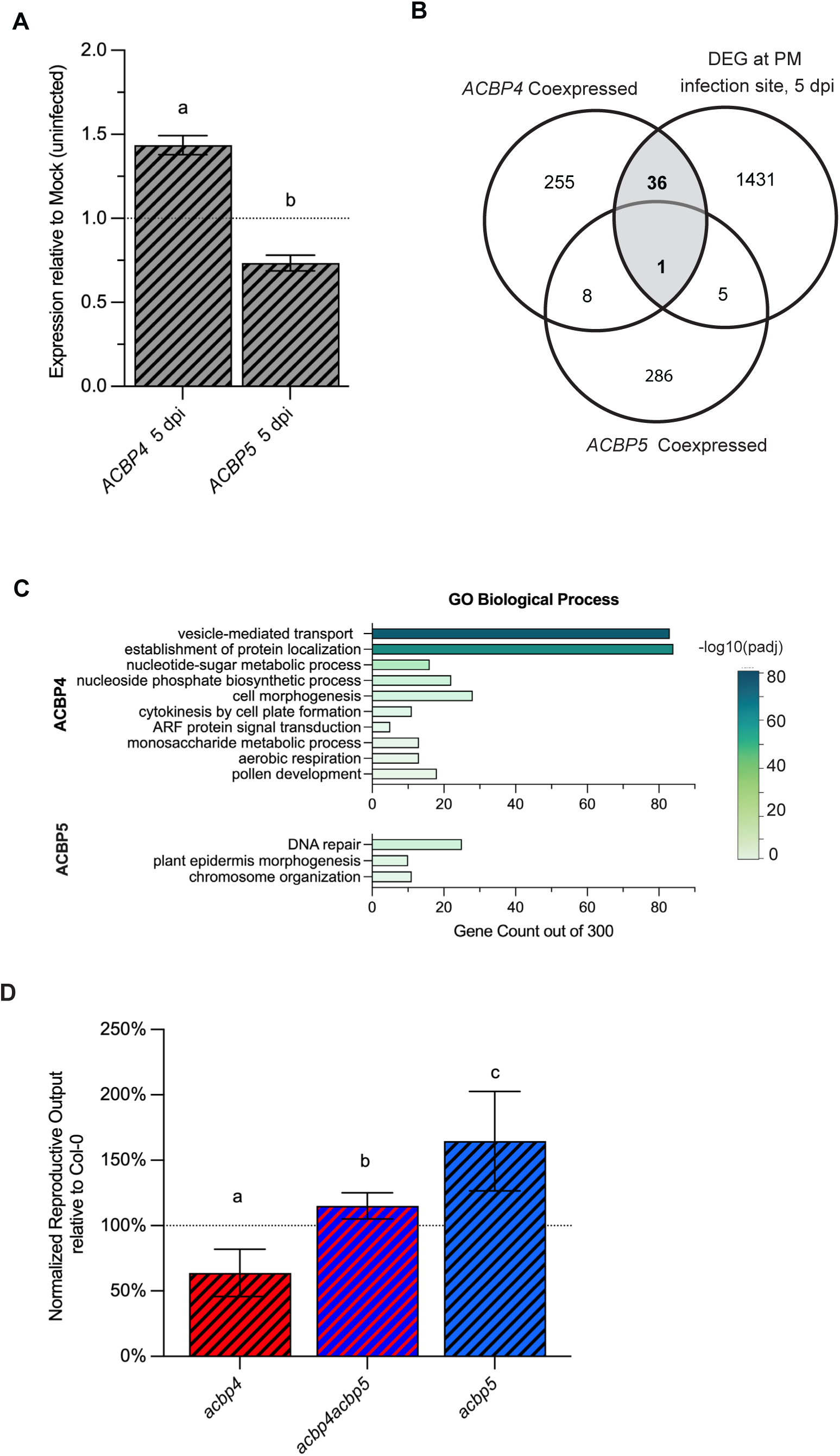
Class IV ACBPs perform distinct roles in the context of powdery mildew infection. **A)** Expression ratios of *ACBP4* and *ACBP5* calculated from mean ± SEM of three independent biological samples at 5 dpi compared to mock (uninfected) leaves of wild-type (WT) Col-0 plants. qPCR for *ACBP4* and *ACBP5* was first normalized to housekeeping gene *ACT2*. Statistically significant differences (p_adj_≤ 0.05; ANOVA and Tukey posthoc test) are denoted by different letters. **B)** Overlap of *ABCP4* and *ACBP5* top 300 coexpressed genes obtained from ATTED-II (Obayashi et al. 2022) with differentially expressed genes (DEG; ≥2-fold, p≤ 0.05) at the powdery mildew (PM) infection site (cells isolated using laser microdissection) at 5 dpi (Chandran et al. 2010). Fisher’s exact probability test found only overlap between *ACBP4* coexpressed genes and DEG genes at PM infection site to be significant (shaded area; p ≤0.0001). **C)** Gene Ontology (GO) term enrichment of biological processes for the top 300 coexpressed genes with *ACBP4* or *ACBP5* is shown for GO Parent Categories, with p_adj_ cutoff of E-5, ordered by p_adj_. Coexpressed genesets retrieved from ATTED-II and biological process classification done through g.profiler (Kolberg et. al, 2023). See Supplemental Workbook S1 for gene sets and analysis output. **D)** Powdery mildew reproductive output of Col-0, *acbp4, acbp4acbp5*, and a*cbp5*. Spore counts are relative to WT control plants at 9 dpi; mean ± SEM, n=6. Letters show significance among genotypes using one-way ANOVA with Tukey post-hoc test. Significant differences from WT determined using paired 2-tailed t-test were p ≤ 0.01. Dark gray= wild-type; Red= *acbp4*; Blue =*acbp5*; Stripes indicate powdery mildew infection.

Analysis of the top 300 genes coexpressed with *ACBP4* and *ACBP5*, obtained from ATTED-II (Obayashi et al. 2022), shows minimal intersection (Fig. 2B, Supplemental Workbook S1). Furthermore, only coexpressed *ACBP4* genes share significant overlap (p ≤ 0.0001) with genes differentially expressed at the Gor infection site (isolated using laser microdissection (LMD)) at 5 dpi (Chandran et al. 2010) (Fig. 2B, Supplemental Workbook S1). Finally, analysis of gene ontology (GO) biological process term enrichment for the top 300 coexpressed genes with *ACBP4* or *ACBP5* suggests they participate in distinct biological processes (Fig. 2C, Supplemental Workbook S1). The most enriched GO terms show no overlap, with ACBP4 associated with vesicle trafficking *(adj. p-value* E-76; GO:0016192) and ACBP5 associated with DNA repair (*adj. p-value* E-12; GO:0006281).

To further assess the relationship between Arabidopsis ACBP4 and ACBP5, we crossed them and obtained a homozygous double *acbp4acbp5* mutant. The double mutant supports slightly more spore production than does WT and an intermediate effect compared to the single *acbp4* or *acbp5* mutants for Gor spore production (Fig. 2D). Taken together our data indicates ACBP4 and ACBP5 play unique roles in their impact on powdery mildew proliferation. Therefore, we concentrate further efforts on uncovering the role of ACBP4 in supporting powdery mildew proliferation.

### ACBP4 is responsible for reduced powdery mildew proliferation

In addition to our quantitative endpoint assessment of powdery mildew proliferation (spore production) for *acbp4* (SALK_040164C, *acbp4-1*) above, qualitative assessments consistently show reduced powdery mildew proliferation with representative images shown in Figure 3A-B. Wild-type plants support more visible white powdery mildew (Fig. 3A top) and show increased density of asexual reproductive structures (conidiophores) (Fig 3A bottom) compared to *acbp4-1* at 7 dpi. Furthermore, conidiophores produced per Gor colony at 5 dpi are reduced on *acbp4-1* compared to WT at 5 dpi (Fig. 3B), consistent with our quantitative analyses showing decreased spore production on *acbp4-1* (Fig. 1B).

**Figure 3:**
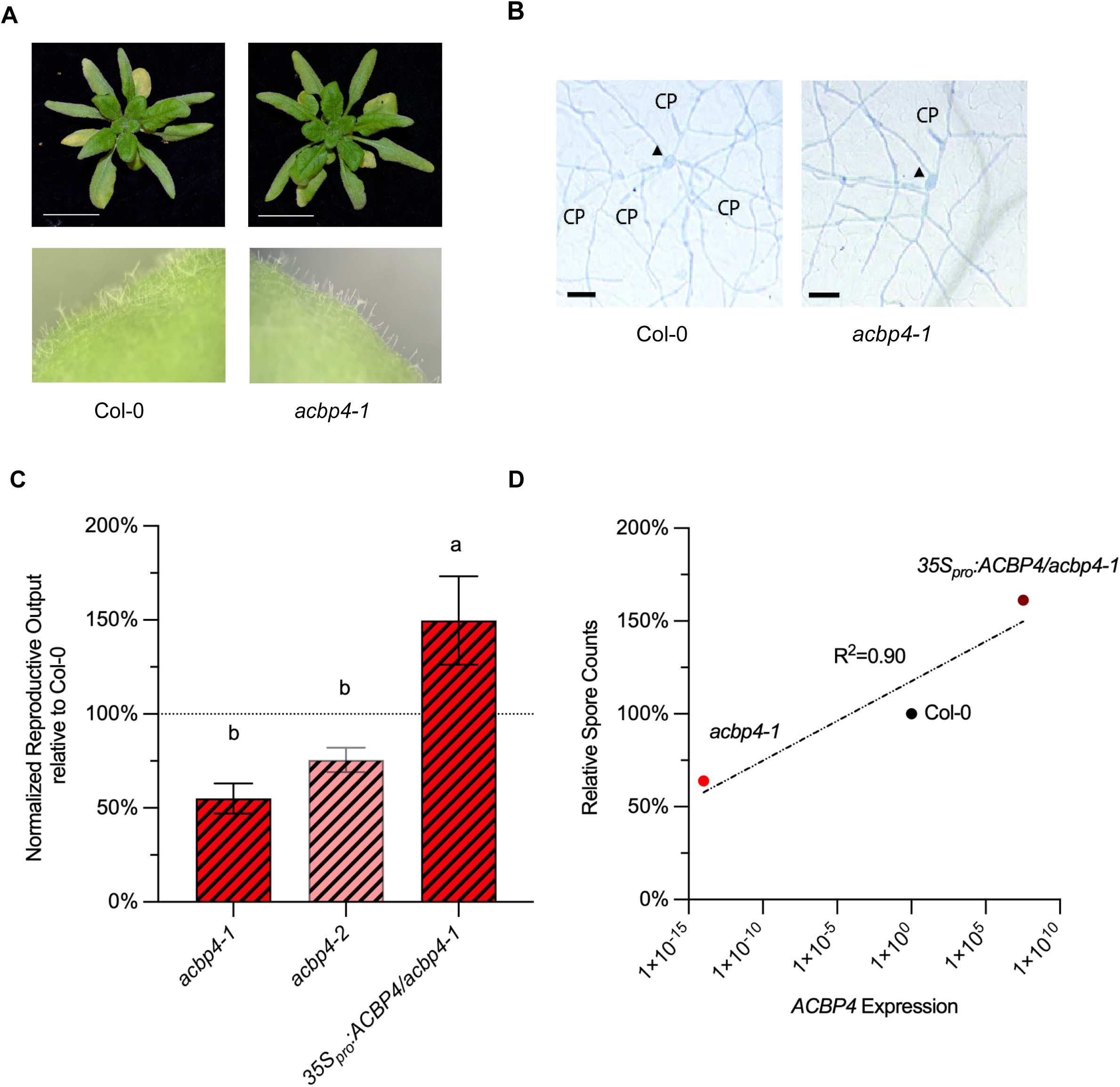
Expression of ACBP4 is linked to powdery mildew growth and reproduction. **A)** Visual disease comparison of representative images of wild-type (WT) Col-0 and *acbp4-1* plants at 7 dpi. Top: whole plant phenotype. Bottom: Close-up of conidiophores on leaf surface. **B)** Colony growth and conidiophores at 5 dpi on WT and *acbp4-1* plants. Scale bar = 50 μm. Arrowhead indicates germinated conidia (spore). CP marks conidiophores. **C)** Powdery mildew reproductive output from WT, independent T-DNA knockout lines, and 3*5S_pro_:ACBP4/acbp4-1* complemented line. Spore counts are relative to WT control plants at 9 dpi; mean ± SEM; n=6. Letters show significance among genotypes using one-way ANOVA with Tukey post-hoc test, p_adj_<0.05. Significant differences from WT determined using paired 2-tailed t-test were p ≤ 0.05. Red = *acbp4*; Stripes= powdery mildew infection. **D)** *ACBP4* expression in Col-0, acbp4-1, and *35S_pro_:ACBP4/acbp4-1* plants, plotted against relative spore counts at 5 dpi. R^2^ determined by linear regression. See Supplementary Figure S1 for details of T-DNA insertion and complemented lines.

To confirm that ACBP4 is responsible for the observed powdery mildew phenotype, we assessed a second independent T-DNA insertion line at the *ACBP4* locus (SALK_114254, *acbp4-2*) and assessed complementation by creating stable transgenics that express *ACBP4* in the *acbp4-1* background (*35S_pro_:ACBP4*/*acbp4*) (Supplementary Figure S1). The independent T-DNA insertion line *acbp4-2* exhibits similar spore reduction as *acbp4-1,* compared to WT (Fig. 3C). Moreover, *35S_pro_:ACBP4*/*acbp4* complements the *acbp4-1* powdery mildew phenotype, with greater *ACBP4* expression and increased spore counts compared to WT plants (Fig. 3C, Supplemental Fig. S1). Therefore, subsequent experiments utilize *acbp4-1* (referred to as *acbp4*). Finally, plotting relative spore production versus *ACBP4* expression for *acbp4*, WT, and *35S_pro_:ACBP4/acbp4* shows them to be highly correlated (*R^2^*= 0.90; Fig. 3D).

### ACBP4 impact on powdery mildew proliferation is not plant defense-associated

To understand the basis of reduced *G. orontii* proliferation on *acbp4* plants, we examined plant defense responses known to limit its growth and reproduction: callose deposition, cell death, and salicylic acid (SA)-mediated defense. Enhanced callose deposition at powdery mildew penetration sites limits powdery mildew colonization (Ellinger et al., 2013). No difference in the intensity of callose staining at penetration sites is observed between WT and *acbp4* (Figure 4A). Plant biotrophs, including powdery mildews, inhibit cell death and are highly impacted by its occurrence (e.g. Wang et al. 2011). Neither constitutive (basal uninfected) nor induced (Gor-infected) cell death is observed in *acbp4* (Fig. 4B).

**Figure 4:**
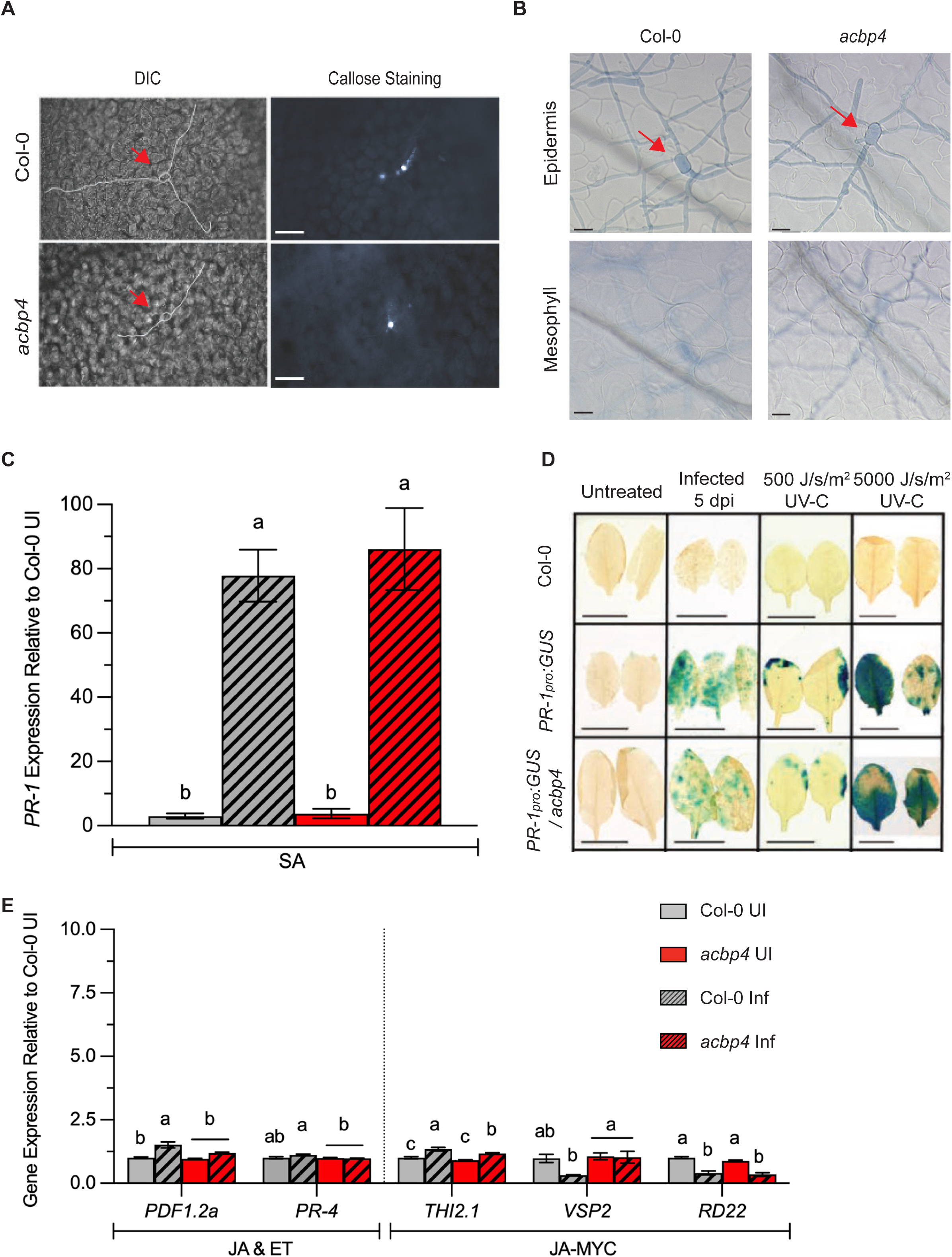
*ACBP4* impact on powdery mildew is not associated with defense. **A)** Callose deposition shown at 2 dpi for Gor colony on Col-0 WT and *acbp4*. Colony growth is outlined in DIC images; callose is visualized by staining leaves with aniline blue. Scale bar = 100 μm. **B)** Trypan blue-stained leaves to assess cell death in epidermal and mesophyll cells at 5 dpi for a Gor colony on Col-0 and *acbp4*. Scale bar = 50 μm. **C)** Gene expression of salicylic acid (SA) marker gene *PR-1* in Col-0 WT and *acbp4* mock-infected (uninfected) and 5 dpi leaves by qPCR, mean ± SEM, n=3. Statistically significant differences (p≤ 0.05; ANOVA and Tukey post-hoc test) are denoted by different letters for ANOVA. **D)** *PR-1_pro_:GUS* reporter signal comparison in *PR-1_pro_:GUS x* Col-0 and *PR-1_pro_:GUS x acbp4*. Conditions are untreated, 5 dpi, low UV-C, and high UV-C treatments. Images are representative of phenotypes from two independent experiments of each treatment comparison; each experiment used 6 plants, of which 2 leaves of matched ages were harvested from each for GUS visualization. Scale bar = 10 mm. **E)** Gene expression of selected markers of jasmonic acid (JA) & ethylene (ET), and JA-MYC defense pathways in Col-0 and *acbp4* mock-infected (uninfected) and 5 dpi leaves, as in **C)**. Statistically significant differences (p≤ 0.05; ANOVA and Tukey post-hoc test) are denoted by different letters for ANOVA, performed on each gene set. Dark gray= WT leaves; Red= *acbp4* leaves; Stripes=infected leaves; No stripes= uninfected leaves.

SA accumulation and defense increase during *G. orontii* infection and limit the extent of infection (Dewdney et al. 2000; Wildermuth et al. 2001; Kuhn et al. 2017). No difference in induced SA defense as assessed by expression of the marker gene *PATHOGENESIS RELATED-1* (*PR-1*) is seen for *acbp4* compared to WT (Fig. 4C). To examine alterations in the pattern of accumulated *PR-1* expression, we crossed a *PR-1_pro_*:GUS reporter line to *acbp4* plants and looked at GUS activity in leaves without and after three SA-induction treatments. There is no difference in GUS activity in the *acbp4* background compared to WT with Gor infection, moderate or high UV-C exposure (Fig. 4D). Taken together, these results indicate that the reduced powdery mildew proliferation observed on *acbp4* is not associated with plant defense responses known to limit powdery mildew growth and reproduction.

Though quantitative plant defense against powdery mildew is largely controlled by SA, alterations in other plant hormone response pathways can modulate powdery mildew growth and asexual reproduction. This may occur through complex cross-talk and feedback on SA signaling and response, often involving jasmonic acid (JA) and ethylene (ET), or on SA-independent impacts. *ACBP4* can be induced by application of the ET precursor aminocyclopropane carboxylic acid or by JA (e.g. Li et al. 2008), and it can bind transcription factors that mediate JA+ET vs. SA antagonism (e.g. WRKY DNA BINDING PROTEIN 70 (WRKY70; Guo et al. 2024) and JA+ET vs JA-MYC antagonism (e.g. ARABIDOPSIS ETHYLENE-RESPONSIVE BINDING PROTEIN (AtEBP; Li et al., 2008). JA+ET dependent signaling is reflected by transcriptional activation of *PLANT DEFENSIN1.2* (*PDF1.2*) and *PATHOGENESIS-RELATED 4* (*PR-4*) (Pré et al. 2008). Whereas, JA signaling marker genes activated by the β-helix loop helix transcription factor MYC2 include *THIONIN 2.1 (THI2.1)*, *RESPONSIVE TO DESSICATION22 (RD22)*, and *VEGETATIVE STORAGE PROTEIN 2* (*VSP2*) (Dombrecht et al. 2007; Wasternack and Hause, 2013). Overall, *acbp4* plants show almost no difference in expression of JA+ET or JA-MYC marker genes in either UI or infected tissues at 5 dpi, relative to WT (Fig. 4E). The most notable difference is the maintenance of *VSP2* expression in 5 dpi *acbp4* plants. *VSP2* expression is reduced 3-fold in WT plants in response to Gor infection but is not reduced in *acbp4*. Reduced *VSP2* expression in response to powdery mildew in WT plants is consistent with JA-MYC versus JA+ET antagonism that is slightly modulated by ACBP4. However, the observed difference in expression is very minor and not all marker genes in a category show a corresponding alteration in *acbp4* (e.g. *RD22* in JA-MYC category, Fig. 4E).

Our results indicate that altered defense in *acbp4* does not account for the powdery mildew asexual reproduction phenotype. This is further supported by GO analysis of the top 300 genes coexpressed with *ACBP4* as no defense-associated pathways show highly significant enrichment (Fig. 2C, Supplemental Workbook S1).

### Leaf lipid profiling reveals basal very long chain FAs differ in *acbp4* compared to wild-type

ACBP4 is a cytosolic lipid trafficking protein that preferentially binds FA acyl-CoAs trafficked from the chloroplast to the ER. It prefers 18:0 acyl-CoA as quantitatively assessed using isothermal titration calorimetry (Hsiao et al. 2014), but can also bind the more predominant 18:1 and 16:0 acyl-CoAs, as well as phosphatidylcholine (PC) (Hsiao et al. 2014; Xiao et al. 2008; Xiao et al. 2009). Therefore, ACBP4 has been proposed to traffic FAs produced in the chloroplast to the ER for elongation and synthesis of membrane lipids. This function of ACBP4 could be particularly relevant in support of the specific lipid demands of the powdery mildew during asexual reproduction.

Lipids from mock uninfected and 12 dpi leaves from WT and *acbp4* plants were isolated and analyzed for phospholipids and galactolipids via electrospray ionization with mass spectrometry (ESI-MS) and FA composition via FA methyl ester (FAME) analysis with gas chromatography – flame ionization detection. During tissue harvest, spores growing on the infected leaves were also isolated and prepared for lipid analysis to assess whether alterations in host lipids are reflected in the spore composition.

Summed phospholipid abundance is similar in WT and *acbp4*, with a decrease of ∼20% in response to infection in both genotypes (Figure 5A, Supplemental Workbook S2). However, there is no difference in summed phospholipid abundance by class for *acbp4* compared to WT in infected or uninfected leaf tissue (Fig. 5B). Similarly, Guo et al. (2019) found no difference in phospholipid or FA abundance or composition in *acbp4* compared to WT, excepting a slight increase in C18:3 at the expense of very long chain FAs (Guo et al. 2019). Thylakoid membrane associated lipids, monogalactosyldiacylglycerol (MGDG) and phosphatidylglycerol (PG), decrease for both WT and *acbp4* in response to Gor infection (Fig. 5B), consistent with our previous report for the WT response to Gor infection (Xue et al. 2025). However, in contrast to Xia et al. (2012) who report decreased thylakoid membrane-associated lipids in *acbp4* compared to WT, we find no statistically significant differences (Fig. 5B).

**Figure 5:**
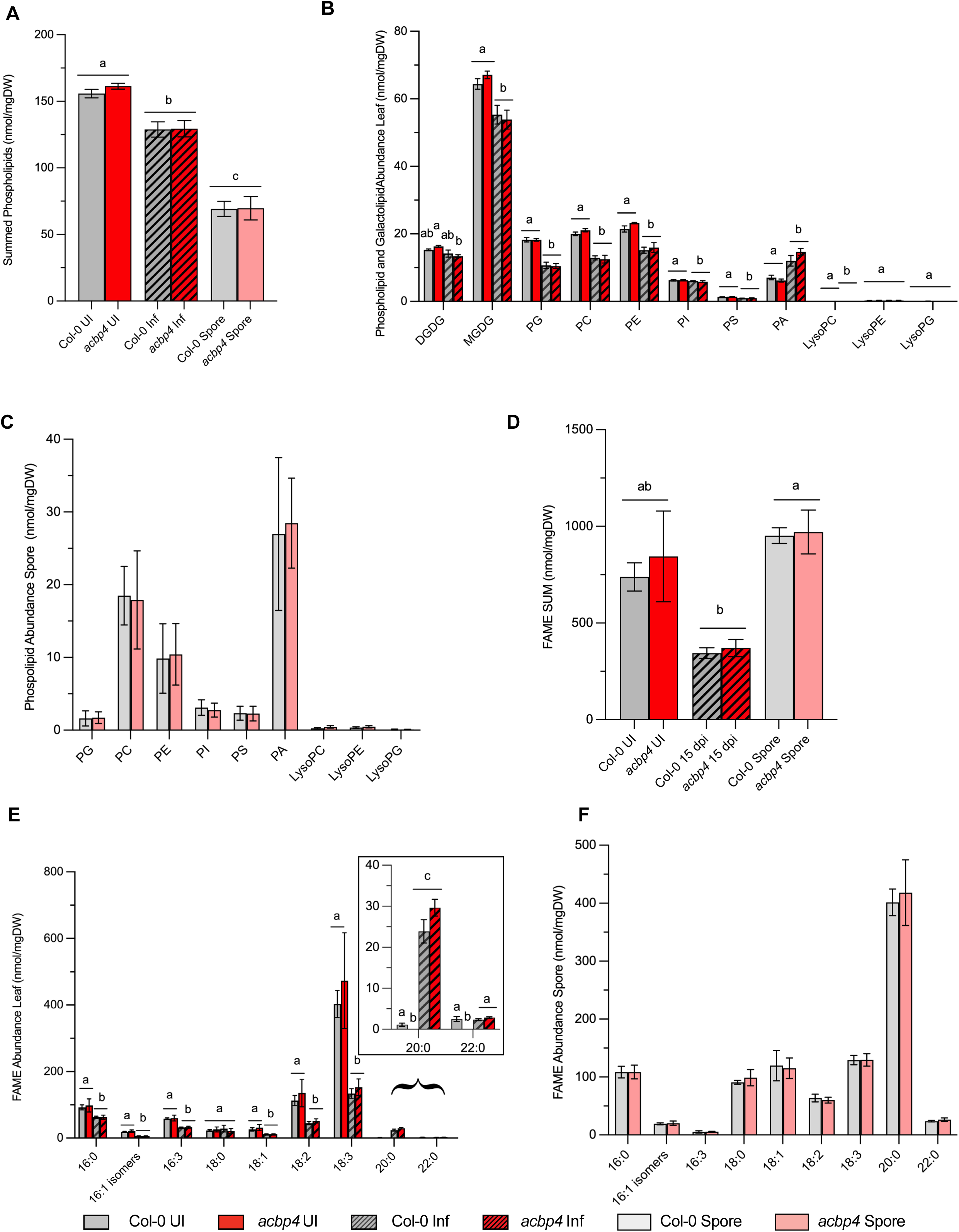
Lipid abundance and acyl chain profiling of Col-0 and *acbp4*. **A)** Summed abundance of phospholipid extracts from uninfected (UI) leaves, infected leaves, and spore tissue from Col-0 and *acbp4* plants, n = 5, mean ± SEM. Statistically significant differences between tissue abundance are denoted by different letters (one-way ANOVA and Tukey post hoc tests,* p ≤ 0.05). **B)** Abundance of detected phospholipids and galactolipids from total lipid extracts from UI and 12 dpi leaves harvested from Col-0 and *acbp4* plants. **C)** Abundance of detected phospholipids from total lipid extracts from spores harvested from infected Col-0 and *acbp4* plants at 12 dpi. **D)** FAME summed abundance in UI and 12 dpi leaves and spores harvested from the Col-0 and *acbp4* plants, mean± SEM, n=5. Statistically significant differences between tissue abundance are denoted by different letters (tested by ANOVA and Tukey post hoc tests, p ≤ 0.05). **E)** Leaf abundance of detected FAMEs by species. Statistically significant differences between genotypes and tissue for each species are denoted by different letters (tested by ANOVA and Tukey post hoc tests, p ≤ 0.05). **F)** Spore abundance of detected FAMEs by species. Differences by genotype were not significant (tested by ANOVA and Tukey post hoc tests, p ≤ 0.05). Dark gray= WT leaves; Red= *acbp4* leaves; Stripes=infected leaves; No stripes= uninfected leaves; Light gray= spores harvested from WT leaves; Light red= spores harvested from *acbp4* leaves.

Total FA abundance is also comparable in WT and *acbp4*, with a decrease of ∼60% with infection in both WT and *acbp4* leaves (Fig. 5D). The decrease in leaf FA abundance with infection is mainly attributable to changes in C18:3 and C18:2 species, which decrease by ∼60% in both WT and *acbp4* leaves at 12 dpi compared to uninfected basal levels (Fig. 5E), consistent with our previous findings for leaves of Gor-infected WT (Xue et al. 2025). By contrast, though very long chain FAs (VLCFAs; >C18) comprise a small proportion of the FA pool, C20:0 increases more than 10-fold in WT plants with infection. The increase of VLCFAs during PM infection was also observed by Xue et al. (2025); C20:0 VLCFAs increase ∼20-fold in infected leaves at 12 dpi. Strikingly, no VLCFAs (C20:0 or C22:0) are detected in uninfected *acbp4* leaves; however, with infection, these VLCFAs are induced to WT levels (Fig. 5E). Therefore, although basal leaf VLCFAs are not detected in *acbp4*, *acbp4* does appear to retain the capacity to synthesize VLCFAs in response to a given inducer, though the possibility remains that Gor contributes to VLCFA in infected leaves as the haustorial complex is retained in washed infected leaves.

Abundance and composition of phospholipids and FAs are not altered for spores produced on *acbp4* compared to WT (Fig. 5A,C-D,F). Unlike leaves, Gor spores are dominated by VLCFAs, with 4-fold more C20:0 than any other FA species (Fig. 5F), consistent with our previous results (Xue et al. 2025). By contrast, basal leaf C20:0 comprises only ∼1% of leaf FAs, though C:20 increases dramatically with infection (to ∼20% in infected leaves) at the whole leaf level (Xue et al. 2025). The localized increase in VLCFAs (e.g. C20:0) at the infection site where lipids are acquired by the fungus may be even greater as we see a 11-fold increase in TAGs at the infection site with a 3.5-fold increase at the whole leaf level (Xue et al. 2025). Given the lack of difference between Gor-infected leaf lipids at 12 dpi for *acbp4* versus WT, it is not surprising that spores developed on those leaves also show no difference in lipid profile. The only significant difference in profiled lipids between *acbp4* and WT is a reduction in basal leaf VLCFA C:20 and C:22 content.

### *acbp4* has reduced leaf basal mesophyll cell ploidy resulting in decreased final ploidy of mesophyll cells underlying the fungal haustoria complex

Gor induces localized host mesophyll cell endoreduplication (2-4 cycles) at the infection site (Chandran et al. 2010; Chandran et al. 2013), which is associated with increased metabolic capacity and flux to lipids (Lee et al. 2024; Xue et al. 2025). The DNA content (ploidy) of leaf mesophyll cells directly underlying the epidermal cell containing the Gor hausorial complex at 5 dpi is highly correlated with the extent of Gor asexual reproduction (Chandran et al. 2010; Chandran et al. 2013). This final ploidy of mesophyll cells underlying the feeding structure in Gor-infected tissue can be altered by changes in the basal cell ploidy or the extent of the Gor-induced endocycles (Chandran et al. 2013).

Using confocal microscopy with 4’,6-diamidino-2-phenylindole (DAPI) DNA staining and 3D reconstruction of Z-stacks, we quantified the ploidy of the three mesophyll cells underlying the epidermal cell containing the haustorium at 5 dpi and of similarly positioned leaf mesophyll cells of mock uninfected plants grown and maintained in parallel for *acbp4* and WT, as in Chandran et al. (2010; 2013). In both uninfected and infected leaves, the localized ploidy distribution for *acbp4* is shifted to the left (lower ploidy) compared to WT (Figs. 6A-B, Fig. S2A-B). The ploidy distribution is represented by the ploidy index, with ploidy index = (%4C nuclei × 1) + (%8C nuclei × 2) + (%16C nuclei × 3) + (%32C nuclei × 4) + (%64C nuclei × 5) as in Chandran et al. (2010; 2013). As clearly shown by comparison of ploidy indices, the decreased final mesophyll cell ploidy at the infection site at 5 dpi of *acbp4* is not attributed to reduced Gor-induction of endoreduplication, but instead to lower initial basal ploidy (Fig. 6C). Representative confocal Z-stack images of conidia on the surface of infected leaves and the three mesophyll cells underlying the infection site are shown in Fig. 6D. The WT Col-0 ploidy indices for basal and 5 dpi are consistent with our previous studies (Chandran et al. 2010; 2013). Furthermore, inclusion of the *acbp4* data with previous findings for other mutants with altered final mesophyll cell ploidy at the infection site (Chandran et al. 2013) shows it maintains a high correlative value for normalized asexual reproductive output with infection site-specific mesophyll cell ploidy at 5 dpi (*R^2^* = 0.76; Supplemental Figure S2). This suggests ACBP4 limits the extent of powdery mildew asexual reproduction through its impact on host susceptibility, and specifically basal (developmentally programmed) endoreduplication.

**Figure 6.**
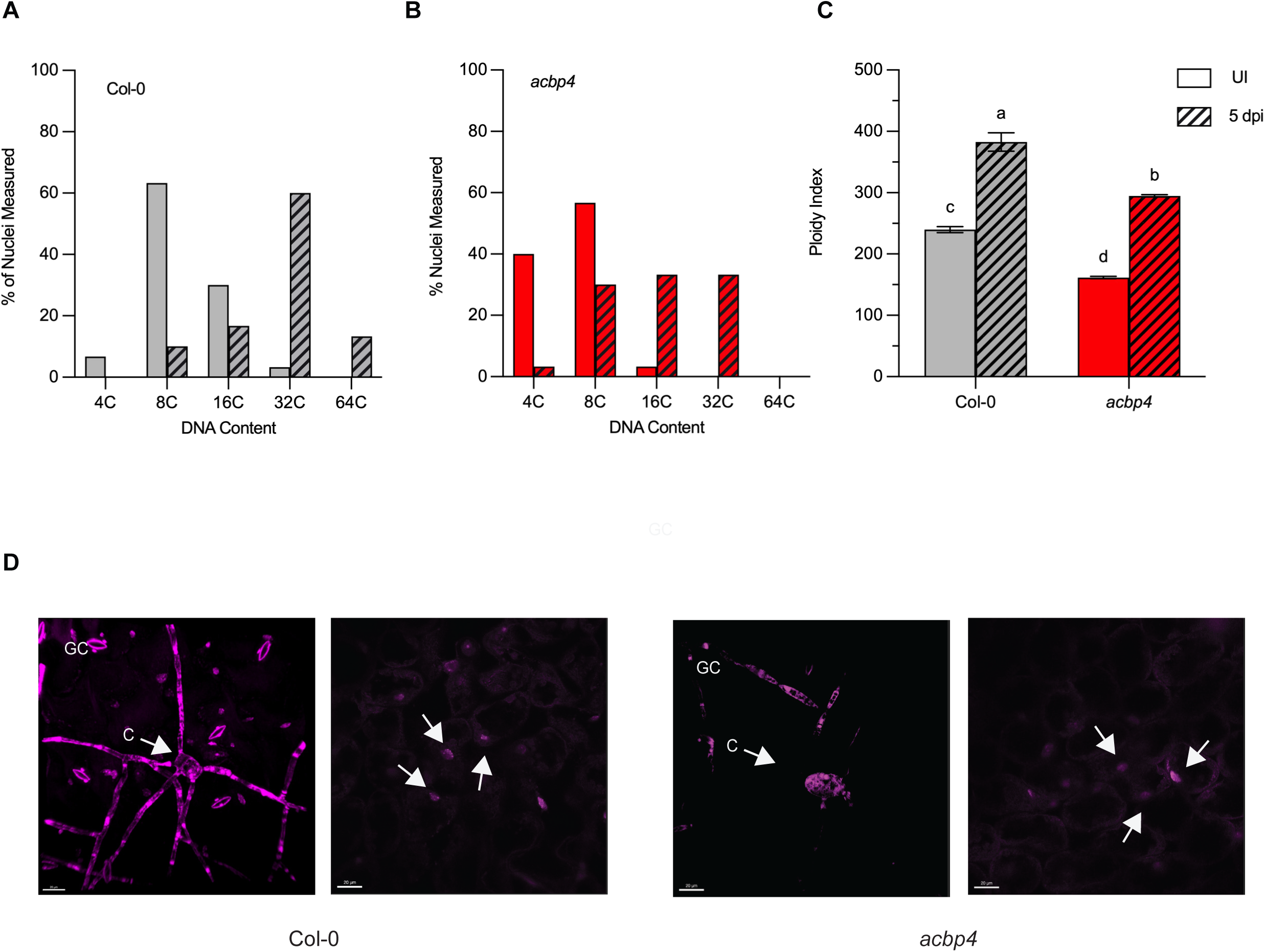
Basal ploidy and ploidy of leaf mesophyll cells at the powdery mildew infection site is altered in *acbp4* compared to wild-type plants. **A)** Ploidy distribution of the three mesophyll cells underlying the haustoria is altered in Col-0 uninfected tissue as compared to powdery mildew infected tissue at 5 dpi as a percent of total nuclei measured. n ≥ 30 nuclei. One representative experiment is shown, the other is shown in Supplementary Figure S2A. Both independent experiments yielded similar results. **B)** Ploidy distribution of the three mesophyll cells underlying the haustoria is altered in *acbp4* uninfected (UI) tissue as compared to infected tissue at 5 dpi as a percent of total nuclei measured. n ≥ 30 nuclei. One representative experiment is shown, the other is shown in Supplementary Figure S2B. Both independent experiments yielded similar results. **C)** Ploidy index of uninfected and infected WT and *acbp4* plants calculated by Ploidy index = (%4C nuclei × 1) + (% 8C nuclei × 2) + (%16C nuclei × 3) + (%32C nuclei × 4) + (%64C nuclei × 5). Statistical differences were calculated by ANOVA and Tukey post-hoc test (p≤0.05). **D)** Representative confocal Z-stack images of leaf surface (left) showing DAPI-stained conidia (C), and stomatal guard cell (GC), and underlying Z-stack image (right) showing three mesophyll cells (M) underlying the haustoria quantified for ploidy. GC are used for normalization of ploidy to 2C. Dark gray= wild type Col-0; Red= *acbp4*; Stripes= powdery mildew infected; No stripes= uninfected.

### Ploidy-associated impacts are also observed in the leaf epidermis and in dark grown hypocotyls of *acbp4* compared to WT

Mature *acbp4* plants look similar in size to WT (Fig. 3A) and the rosette diameter of *acbp4* plants does not differ from WT (Fig. 7A). However, there is an increased number of stomata per area (stomatal density) on the adaxial epidermis of mature leaves of *acbp4* relative to WT plants (Fig. 7B). The stomatal index (number of stomata divided by number of epidermal pavement cells) is unchanged in *acbp4* relative to WT (Fig. 7C), suggesting the epidermal pavement cells in *acbp4* are smaller than WT. Indeed, *acbp4* plant epidermal pavement cells are on average ∼50% smaller than those of WT leaves (Fig. 7D-E). Stomatal density in Arabidopsis has been shown to be inversely correlated with epidermal pavement cell size and ploidy (Li et al. 2020; Melargno et al. 1993). Therefore, our results are consistent with reduced endoreduplication (basal cell ploidy) in these adaxial leaf epidermal cells.

**Figure 7:**
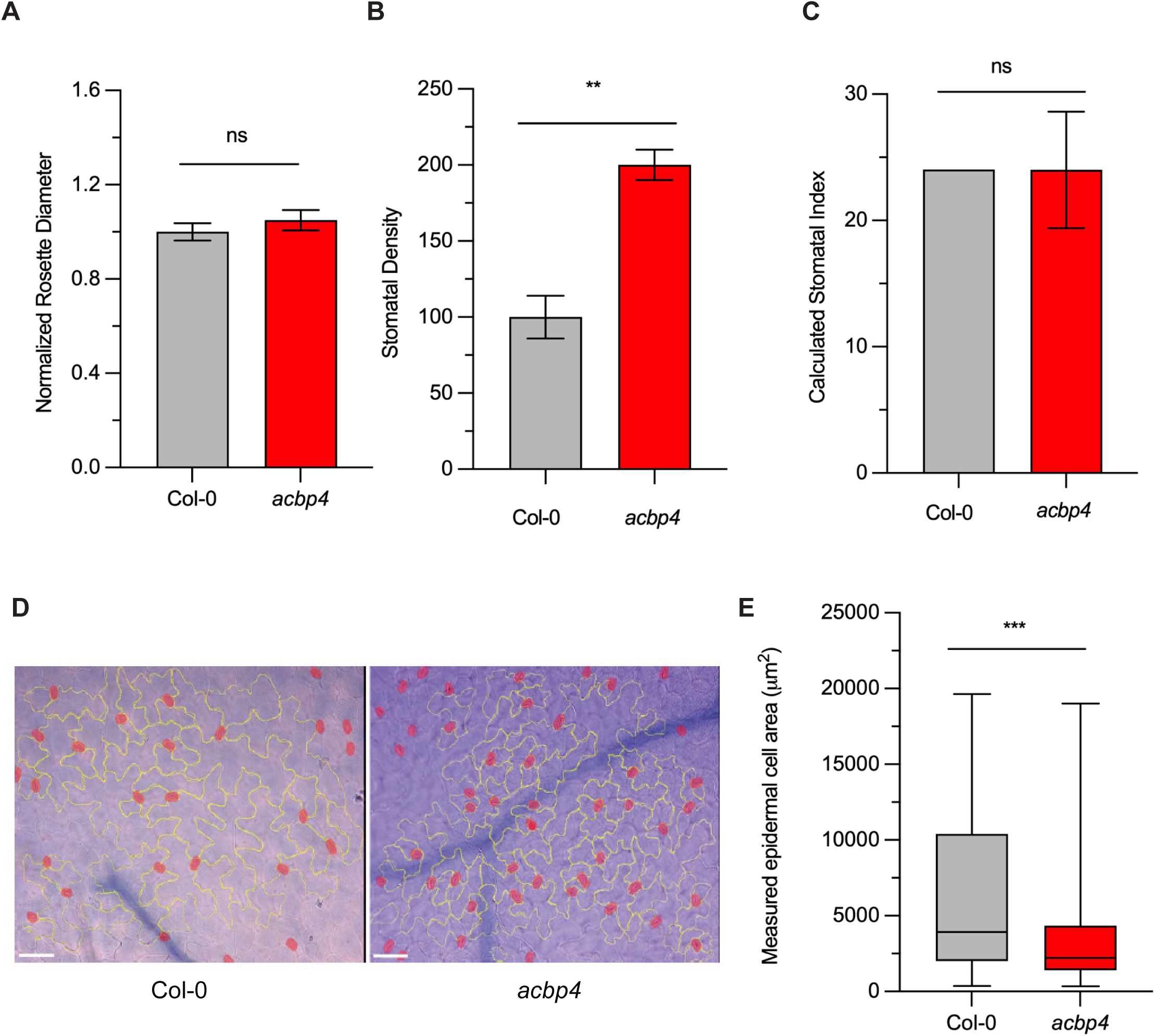
Comparison of rosette size, stomatal density, index, and epidermal cell size. **A)** Rosette diameter at 3.5 weeks of Col-0 and *acbp4* plants n = 45 ± SEM. No significant difference detected by 2-tailed unpaired T-test (* p ≤ 0.05). **B)** Adaxial leaf surface stomatal density at 3.5 weeks (stomata/50µm^2^), n = 3 ±SEM. Significant difference detected by 2-tailed unpaired t-test (* p ≤ 0.05). **C)** Adaxial leaf surface stomatal index at 3.5 weeks, calculated as (number of stomata)/(total number of epidermal cells)*100, as described in Royer, 2001). n = 3, ± SEM. No significant difference detected by 2-tailed unpaired T-test (p ≤ 0.05). **D)** Representative images of adaxial leaf surface. Epidermal pavement cells are outlined in yellow. Stomata are highlighted in red. Bar = 80µm. **E)** Measured epidermal pavement cell areas. Data from 3 independent leaf sets of counts. Dark gray= Col-0 WT; Red=*acbp4*.

To further investigate whether ACBP4 contributes to developmentally programmed endoreduplication, we examined hypocotyl elongation in the dark, a process that is driven by endoreduplication-associated cell expansion (Gendreau et al. 1998; Narukawa et al. 2015). We find that hypocotyl length is decreased in *acbp4* compared to WT (Fig. 8A-B), as is epidermal cell length (Fig. 8C) and ploidy (Fig. 8D). The most dramatic differences in cell ploidy of dark-grown elongating hypocotyls for *acbp4* compared to WT are the increased number of cells in the 2C ploidy class and decreased number of cells in the 16C ploidy class (Fig. 8D), consistent with decreased endoreduplication events in *acbp4* compared to WT.

**Figure 8:**
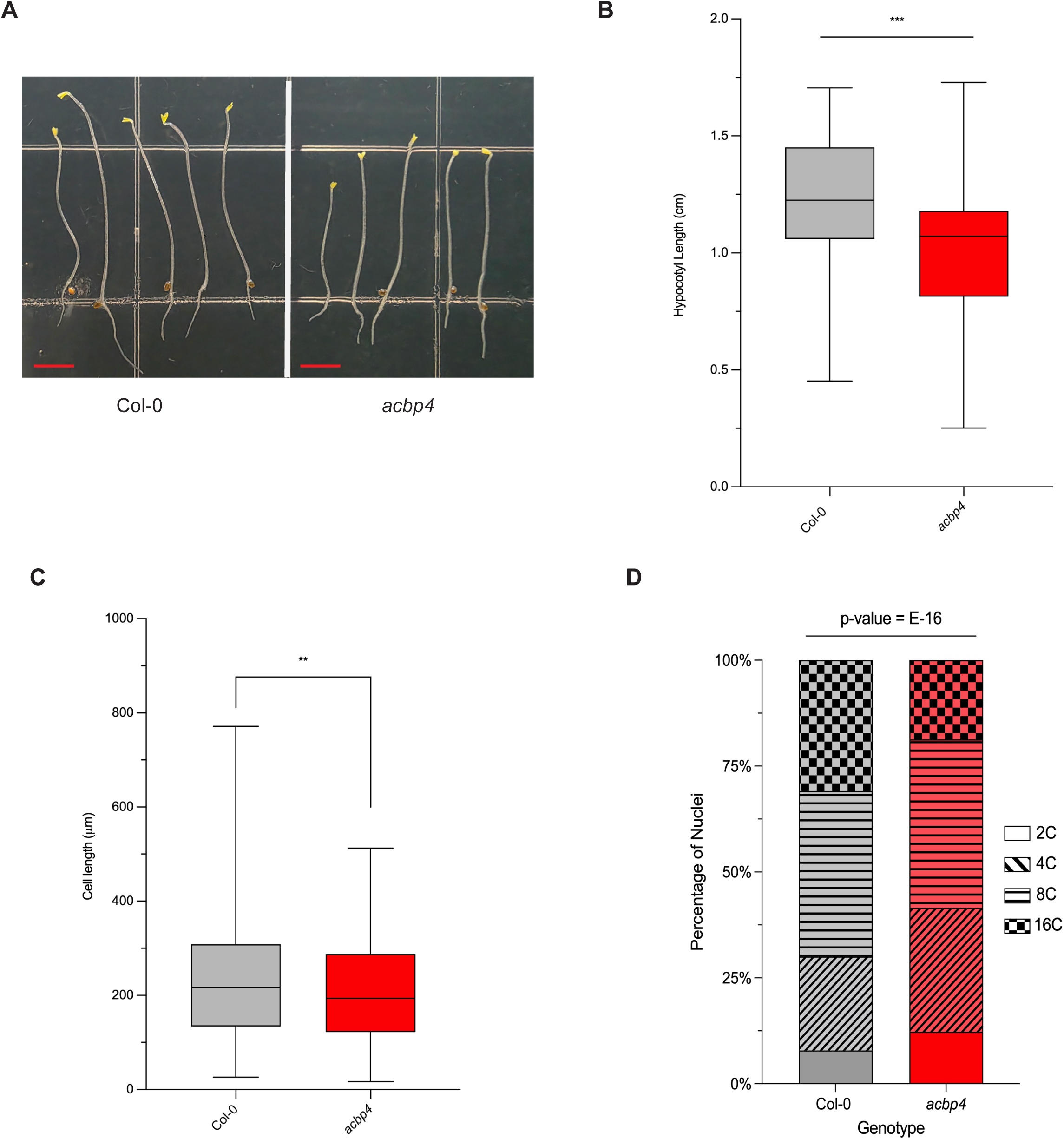
*acbp4* hypocotyls exhibit reduced growth, ploidy, and cell length in dark grown hypocotyls. **A)** Representative image of Col–0 (left) and *acbp4* (right) dark grown hypocotyls at 4 days. Scale bar = 0.5 cm. **B)** Hypocotyl lengths (cm) were measured for 4 day old Col-0 and *acbp4* seedlings grown on medium containing DMSO. In the box plots, the center line represents the median, the box limits indicate the upper and lower quartiles, and the whiskers extend to the highest and lowest values within 1.5 times the interquartile range. Sample sizes: Col-0 DMSO (n = 60), *acbp4* DMSO (n = 57) .One representative experiment is shown, the other is shown in Supplementary Figure S3. Both independent experiments yielded similar results. Significance tested by two-tailed t-test followed by Tukey’s HSD post-hoc test (*p ≤ 0.05, **p ≤0.01 ***p ≤0.001). **C)** Hypocotyl epidermal cell lengths (μm) of 4 day old Col-0 and *acbp4* seedlings. Sample sizes: Col-0 (n = 8 seedlings, 352 total cells), *acbp4* (n = 7 seedlings, 341 total cells). Significance was tested by two-tailed t-test (*p ≤ 0.05, **p ≤ 0.01 ***p ≤ 0.001). **D)** Distribution of nuclear ploidy levels in hypocotyl tissues of 4-day-old dark-grown Col-0 and *acbp4* seedlings, displayed as stacked bars representing the percentage of nuclei in each ploidy class (2C, 4C, 8C, and 16C). Sample sizes: Col-0 (n = 6 seedlings, 1662 total nuclei), *acbp4* (n = 6 seedlings, 1547 total nuclei). Significance in ploidy distribution between genotypes was tested by Pearson’s Chi-square test of independence (df=3; p-value = E-16). Dark gray= Col-0 WT; Red= *acbp4*.

Taken together, we show reduced basal leaf mesophyll cell ploidy (Fig. 6C), reduced epidermal pavement cell size and increased stomatal density (Fig. 7B,D-E) consistent with reduced leaf epidermal pavement cell ploidy, and reduced hypocotyl length, epidermal cell length and ploidy in dark-elongated hypocotyls (Fig. 8). These findings indicate that ACBP4 plays a role facilitating developmentally programmed endoreduplication.

## DISCUSSION

### Powdery mildew infection phenotype highlights distinct functions of ACBP4 and ACBP5

AtACBP4 and AtACBP5 are highly similar Arabidopsis Class IV acyl-CoA binding proteins. In addition to an N-terminal acyl-CoA binding domain, they contain kelch repeat propeller domains, and a C-terminal coiled-coil domain (Fig. 1A). Both are soluble, largely cytosolic proteins with shared acyl-CoA preferences by lipidex-binding analyses (Xiao et al. 2008, 2009). However, quantitative isothermal titration calorimetry (ITC) assays show AtACBP4 has >10-fold greater binding affinity for 18:0 than for any other acyl-CoA tested; AtACBP5 Kds are similar to those of AtACBP4 with this exception (Hsiao et al. 2014).

Previous studies indicate complementary functional overlap between Arabidopsis ACBP4 and ACBP5 in floral lipid metabolism (Ye et al. 2017), seed development (Guo et al. 2019) and response to hypoxia (Guo et al. 2024). However, utilizing the *A. thaliana - G. orontii* pathosystem, we show these proteins have divergent functionalities, with knockouts exhibiting opposite impacts on powdery mildew spore production (Fig. 1B) and the *acbp4acbp5* double mutant exhibiting an intermediate spore production phenotype (Fig. 2D).

A strong transcriptional Arabidopsis response is observed with Gor infection at 5+ dpi and it has facilitated the identification of genes and processes of importance for the support of Gor asexual reproduction on Arabidopsis (Chandran et al. 2009, Chandran et al. 2010; Lee et al. 2024). Only *ACBP4*, not *ACBP5,* expression at 5 dpi is significantly increased during PM infection (Fig. 2A). Moreover, only the top 300 coexpressed genes with *ACBP4*, not *ACBP5*, have a statistically significant overlap (37/300) with those differentially expressed at the powdery mildew infection site at 5 dpi (Fig. 2C). Gene Ontology biological process term enrichment for the top 300 coexpressed genes with *ACBP4* or *ACBP5* further supports unique functional roles for these two proteins (Fig. 2C). ACBP4 is strongly associated with vesicle-mediated transport (*adj. p-value*: E-76), while ACBP5 is associated with DNA repair (*adj. p-value* E-12).

### ACBP4 function in promoting powdery mildew spore production is associated with its impact on mesophyll cell ploidy at the infection site, not plant defense

Quantitative reduction of powdery mildew spore production on Arabidopsis mutants has been largely attributed to enhanced plant defense or compromised host susceptibility. We first examined plant defenses known to most dramatically impact powdery mildew growth and reproduction including callose deposition, cell death, and SA-mediated defense. None were altered in *acbp4* compared to WT (Fig. 4). In addition, as ACBP4 can bind the AtWRKY70 transcription factor that mediates JA-ET vs. SA antagonism (Guo et al. 2024) and AtEBP involved in JA+ET vs. JA-MYC antagonism (Li et al. 2008), we also assessed the transcriptional response of markers in these pathways (Fig. 4). Reduced *VSP2* expression (3-fold) and 1.6-fold enhanced expression of *PDF1.2* in infected WT plants could potentially be attributed to EBP, or even WRKY70, context-dependent function via repression of JA-MYC genes (e.g. *VSP2*) and activation of ET+JA genes (e.g. *PDF1.2*). Therefore, it is possible that a minor impact of ACBP4 in the powdery mildew interaction functions through disruption of ACBP4-WRKY70 or ACBP4-EBP interactions. However, the observed differences in expression are minimal with no concerted change in expression for marker genes in a given category of response. Taken together, our results indicate that alterations in plant defense do not drive the *acbp4* reduced powdery mildew asexual reproduction phenotype.

Compromised host susceptibility that quantitatively impacts *G. orontii* spore production includes alterations in host cell ploidy underlying the fungal feeding site (Chandran et al. 2010, 2013) and host lipid metabolism (Lee et al. 2024; Xue et al. 2025). Increased host cell ploidy underlying the fungal feeding site is associated with enhanced metabolic capacity and the extent of powdery mildew asexual reproduction is highly correlated with this host mesophyll cell ploidy at 5 dpi, consistent with its function as a host susceptibility factor (Chandran et al. 2013). Concurrently there is a localized shift in host lipid metabolism to increase flux to lipids and accumulate specific lipids that support powdery mildew spore production (Lee et al. 2024; Xue et al. 2025). For *acbp4*, basal (uninfected) mesophyll cells exhibit lower cell ploidy than WT (Fig. 6). Therefore, although localized Gor-induced host cell endoreduplication still occurs, the final mesophyll cell ploidy at 5 dpi in *acbp4* is lower than it is for WT. The *acbp4* result corroborates with our previously reported (Chandran et al. 2013) Gor asexual reproduction - mesophyll cell ploidy correlation (R^2^=0.75, Supplemental Figure S2) suggesting the reduced spore production on *acbp4* is largely due to the reduction in basal mesophyll cell ploidy. A similar situation is observed for mutants in *POWDERY MILDEW RESISTANT 6* (*PMR6*) and *PLANT UBIQUITIN X BOX 2* (*PUX2*) which have reduced Gor asexual reproduction and basal mesophyll cell ploidy (Chandran et al. 2013). Furthermore, the reduced Gor growth and reproduction phenotype on *acbp4* appears to be specific to infection with powdery mildew, a fungal obligate biotroph, as Xia et al. (2012) found *acbp4* to be more susceptible to two fungal necrotrophs and the bacterial hemibiotroph *Pseudomonas syringae* pv. *tomato* (Pst). Similarly, mutants in *PMR6* exhibit divergent phenotypes. *pmr6* is equally susceptible to Pst (Vogel et al. 2002) and more susceptible to fungal necrotrophic pathogens than wild-type (Chiniquy et al. 2019). Although *pux2* has not been tested for altered susceptibility to these pathogens, it was identified as a susceptibility factor in tobacco, facilitating infection by the haustoria-forming hemibiotroph oomycete *Phytophthora nicotianae* (Meng et al. 2021), as it does for Gor on Arabidopsis (Chandran et al. 2013).

### ACBP4 functions in developmental endopolyploidy

In addition to reduced basal mesophyll cell ploidy in mature Arabidopsis leaves (Fig. 6), we also observe reduced epidermal pavement cell size and increased stomatal density (Fig. 7), which correlate with lower epidermal pavement cell ploidy (Li et al. 2020; Melargno et al. 1993). As post-germination hypocotyl growth in the dark is attributed to cell expansion, driven by endoreduplication (Narukawa et al., 2015; Gendreau et al. 1998), *A. thaliana* hypocotyl elongation in the dark can serve as a useful system for the identification of factors impacting developmental endopolyploidy. For example, mutants in *HYP6* with reduced hypocotyl elongation in the dark identified DNA topoisomerase VI as playing a central role in developmental endoreduplication (Sugimoto-Shirasu et al. 2002). We found *acbp4* exhibits reduced hypocotyl length in the dark, with shorter cells of reduced ploidy compared to WT (Fig. 8). The magnitude of impact of *acbp4* on dark-grown hypocotyls (Fig. 8) was reduced compared to its impact on mature leaf epidermal and mesophyll cells (Figs. 6, 7). This differential extent of impact has also been observed for other regulators of developmental endoreduplication, such as SIAMESE (Walker et al. 2000; Churchman et al. 2006), a cyclin-dependent kinase inhibitor that acts in concert with FZR2/CCS52A1, a core activator of the anaphase promoting/cyclosome complex (APC/C), to enable endoreduplication (Kasili et al. 2010; Breuer et al. 2012). Together, our findings provide strong evidence that ACBP4 is a facilitates developmental endopolyploidy.

Transcriptomic evidence further supports a role for ACBP4 in developmental endoreduplication. *CCS52A1* is among the top 300 genes (rank 89) most strongly coexpressed with *ACBP4* (Supplemental Workbook S1). Furthermore, *ACBP4* is the 7th ranked most strongly coexpressed gene with *CCS52A1* (Supplemental Workbook S1). Similar to *acbp4*, *ccs52A1* mutants exhibit reduced mesophyll and epidermal pavement cell size (Larson-Rabin et al. 2009) and support less powdery mildew asexual reproduction (Chandran & Wildermuth, 2016). Another highly coexpressed gene with *ACBP4* is *STOMATAL CYTOKINESIS DEFECTIVE 1* (*SCD1*), a regulator of vesicle trafficking and cytokinesis (Supplemental Workbook S1). Like *acbp4*, Arabidopsis mutants deficient in *SCD1* exhibit reduced leaf epidermal pavement cell size and increased stomatal density (Falbel et al. 2003). In addition, the most highly significant GO enriched biological process term for the top 300 coexpressed genes with *CCS52A1* or *SCD1* is vesicle-mediated transport, with *adj. p-values* of E-41 and E-80, respectively, as it is for *ACBP4* (E-76) (Supplemental Workbook S1). Vesicle trafficking of lipids, carbohydrates, and proteins that synthesize and modify cell membrane and cell wall components is critical to nuclear and cell expansion that accompanies each endoreduplication cycle. Vesicle trafficking is also an essential component of cytokinesis and formation of the cell plate. GO terms enriched for the top 300-coexpressed genes with *ACBP4* also include these processes (e.g. cell growth (E-11) and cytokinesis (E-08)) as well as the trafficked cargo, including cell wall biogenesis (E-7) and nucleotide sugar biosynthesis (E-13) (Supplemental Workbook S1).

Arabidopsis mutants exhibiting phenotypes similar to *acbp4*, such as *pux2* and *pmr6*, can provide additional insight into the interplay of vesicle trafficking, cytokinesis, and endoreduplication. Like Arabidopsis PUX1, PUX2 can act as a specific CDC48A adaptor, facilitating the interaction between the AAA-ATPase chaperone CDC48A that localizes to the cell plate and SYP31, a protein involved in membrane trafficking (Rancour et al. 2004). The top 300 coexpressed genes with *PUX2* are also most highly enriched in vesicle-mediated trafficking (E-14, Supplemental Workbook S1) and PUX2 has unique features not shared by other Arabidopsis PUX proteins that may facilitate a specialized function (Zhang et al. 2026). Mutants in *PMR6*, a pectin lyase-like protein exhibit reduced basal leaf mesophyll cell size and ploidy, epidermal pavement cell size, reduced hypocotyl elongation in the dark, and reduced powdery mildew asexual reproduction (Vogel et al. 2002; Chandran et al. 2013; Li et al. 2020). The decreased pectin and uronic acid content of *pmr6* (Vogel et al. 2002) may explain the impact of *pmr6* on cell size and ploidy as pectin and uronic acid accumulate at the newly formed cell plate during cytokinesis and are also associated with the ability of plant cell walls to expand (Gu and Rasmussen 2022). Together, these findings indicate that specific aspects of vesicle trafficking associated with cytokinesis and cell expansion support developmental endoreduplication. Cytokinesis precedes cell division in plant cells; therefore, genes associated with cytokinesis are generally assumed to support mitosis, not endoreduplication. However, in the right context, some key players associated with cytokinesis (and mitosis) can promote endoreduplication, as shown for MYB3R4, a transcription factor with three MYB domain repeats (Haga et al. 2007; Saito et al. 2015; Chandran et al. 2010). Our analyses suggest the specific alterations in cell plate/cytokinesis-associated components and their trafficking promote endoreduplication during normal development.

### How might ACBP4 function to promote developmentally programmed endoreduplication?

Arabidopsis ACBP4 contains an ABD, kelch propeller domain repeats, associated with protein-protein interactions, and a coiled-coil domain which could support vesicle tethering and membrane fusion (Fig. 1A). Furthermore, ACBP4 contains multiple detected phosphorylation sites (Willems et al. 2024) that could alter specific domain function, protein stability, and/or localization. The combinatorial function of these domains could result in distinct context-dependent functional impacts, such as developmentally programmed endoreduplication.

Acyl-CoA Binding: Our leaf lipid analyses of *acbp4* mutants found no differences in lipid abundance compared to WT, with the exception of VLCFAs 20:0 and 22:0 (Fig. 5). ACBP4 binds 18:0 acyl-CoA with 10-fold higher affinity than other tested acyl-CoAs (Kd = 2.7 uM) and 10-fold higher affinity than the other soluble Arabidopsis ACBPs (Hsiao et al. 2014). Export of 18:0 from the plastid to the ER is typically minor, ∼1-2% of total FA export, but critical to formation of saturated VLCFAs, such as 20:0 and 22:0 in the ER. Therefore, the exceptionally high affinity of ACBP4 for 18:0 acyl-CoA would allow it to function in channeling these specific acyl-CoAs as precursors for saturated VLCFA elongation complexes in the ER. These VLCFAs are critical components of membrane phospholipids and sphingolipids required for vesicle trafficking, membrane expansion, and cell plate formation during cytokinesis (Zheng et al. 2005; Bach et al. 2011; Tang et al. 2020; Zhu et al. 2020). Furthermore, Arabidopsis mutants in the VLCFA elongase *PAS2* show reduced hypocotyl elongation in the dark (Zhu et al. 2020). Though cell ploidy has not been directly assessed in these mutants, their phenotypes support a direct link between VLCFAs, specific cell plate/cytokinesis-associated components, and their trafficking with endoreduplication during normal development.

Protein-protein interactions: In addition to the confirmed interactions of Arabidopsis ACBP4 with the transcription factors AtEBP (Li et al. 2008) and WRKY70 (Guo et al. 2024), a number of cell cycle modulators were identified as putative interactors with ACBP4 by tandem affinity purification (Van Leene et al. 2010). These include INHIBITOR OF CYCLIN-DEPENDENT KINASE 1 (ICK1), which typically limits mitosis, promoting endoreduplication (Zhou et al. 2003; Wang et al. 2000), and CASEIN KINASE-LIKE (CKL) proteins CKL 1, 3, and 4 involved in regulation of cell division and developmental signaling (Tan et al. 2013). ICK1 appears to be strictly localized to the nucleus, while CKLs exhibit both cytoplasmic and nuclear localization. ACBP4 controls WRKY70 nucleo-cytoplasmic localization, dependent on the phosphorylation status of Ser638 of ACBP4 (in the coiled coil domain) and/or binding of 18:1 CoA (Guo et al. 2024). Similarly, ACBP4 could control the localization of ICK1 and/or these CKLs, based on acyl-CoA status and/or phosphorylation. While these interactions and their control remain to be validated and examined, they suggest another possible means by which ACBP4 could integrate lipid metabolic status with endocycle progression.

Together, our analyses suggest a novel role for a plant ACBP, AtACBP4, as an important facilitator of developmental endoreduplication. As discussed above, we propose ACBP4 promotes developmental endoreduplication by coordinating lipid metabolism, vesicle trafficking, and cell-cycle regulatory pathways required for cellular expansion and endoreduplication. Developmental endopolyploidy is associated with enhanced metabolic capacity and typically occurs in plant cells that supply nutrients for rapid growth, expansion, and/or differentiation (e.g. endosperm (Li et al. 2019)) and trichome (Walker et al. 2000)). Numerous agricultural products including oil and fiber crops, tubers, and fleshy fruits and berries depend on developmental endoreduplication. An extreme example is tomato where endoreduplication (up to 512C) occurs as part of tomato fruit development (Chevalier et al. 2014; Bourdon et al. 2010). Understanding novel components of and mechanisms involved in developmentally programmed endoreduplication may allow for increased quality of these products through fine-tuning their development without impacting whole plant ploidy and its trade-offs (e.g. (Pacey et al. 2022; Dominguez Mendez and Studer 2026).

## MATERIALS AND METHODS

### Plant lines, growth, and infection

Mutant list: Seeds for *acbp1* (SALK_206508), *acbp2* (SAIL_690_G01), *acbp3* (SALK_012290), *acbp4-1* (SALK_040164C), *acbp4-2* (SALK_114254), *acbp5* (SALK_134337), *acbp6* (SALK_104339) T-DNA insertion lines in Col-0 background were obtained from Arabidopsis Biological Resource Center (ABRC) at The Ohio State University. All lines, including the *acbp4-1acbp5* double mutant, were genotyped to confirm homozygosity using primers in Supplemental Workbook S3. Supplementary Fig. S1 shows gene map and genotyping results for *acpb4-1* and *acpb4-2*. Wild-type *Arabidopsis thaliana* ecotype Columbia-0 (Col-0) and mutants were grown in SS Metromix200 (Sun Gro) in growth chambers at 22°C with 70% relative humidity under 150μmol m^-2^ s^-1^. After stratification at 4°C, alternating Col-0 and mutant seeds were planted in 6.5 insert boxes (12 plants/box). For whole plant spore count phenotyping, whole boxes were inoculated at 3.5-4 weeks by settling tower with 10-14 dpi conidia from *G. orontii* MGH1 at consistent time of day (Reuber et al. 1998).

### Creation of complemented *acbp4-1* line

ACBP4 (AT3G05420) coding sequence (without stop codon) was cloned from the Arabidopsis Biological Resource Center bacterial clone CD251522 by adding attB recombination sites to both 5’ and 3’ end of sequence using PCR primers listed in Supplementary Workbook S3. Sequence was sequentially cloned into pDONR221 and then pEARLYGATE102 (Earley et al. 2006) plasmid by Gateway cloning (ThermoFisher Scientific). The binary vector was transformed into *Agrobacterium tumefaciens* GV3101. Transformation of Arabidopsis *acbp4-1* was performed using floral dip method (Clough and Bent 1998). *35S_pro_:ACBP4/acbp4-1* line utilized was selected based on recovered *ACBP4* expression and powdery mildew spore production (shown in Fig. 3), and detection of transformed protein by Western Blot (Supplementary Fig. S1).

### Spore tissue collection and counting

Adapted from Weßling and Panstruga (2012) spore counting method. Briefly, at 8-10 dpi, leaves 7-9 from wild-type and mutant plants in a box were harvested. Spores were washed off leaves by vortexing in 15 mL 0.01% Tween-80 for 30 seconds and filtered through 30μm CellTrics filter (Sysmex America) before centrifugation at 4000xg. The resulting spore pellet was resuspended in 200-1000μL water, according to density of infection and use of tissue. For lipid analysis, tissue was immediately frozen and stored until extraction. For spore counting, 3 paired counts of wild type and mutant spore suspensions from a box were performed on a Neubauer improved hemocytometer (Hausser Scientific). Calculation of spores was normalized to the weight of the plant tissue to determine spores/mgFW, and then normalized to wild-type counts. To determine significance, a paired, two-tailed Student’s T-test was performed on counts from at least 5 boxes (p ≤ 0.05).

### Generation of *PR-1_pro_:GUS/acbp4-1 line* and visualization of expression

*Homozygous PR1_pro_:GUS* in Col-0 (as in (Chandran et al. 2010)) was used to pollinate homozygous *acbp4-1*, and selfed. Progeny was screened for the presence of the T-DNA insertion and the *GUS* transgene. F2 progeny were planted in boxes with Col-0 negative control and *PR1_pro_:GUS* positive controls and exposed to treatments. For powdery mildew infection, leaves were harvested at 5dpi after inoculation with light dose (2 leaves). For UV-C treatment, plants exposed to UV (254 nm) in Stratalinker 1800 (Stratagene) for 45 seconds at 500 or 5000 J/s/m^2^ and leaves were harvested 48 hours after UV-C treatment. Leaves were immersed in GUS staining solution (0.1 M NaPO_4_, pH=7, 10 mM EDTA, 0.1% Triton X, 1 mM K_3_[Fe(CN)_6_], 2 mM X-Gluc) and left under vacuum (200 torr) for 15 minutes. Samples were incubated for 24 hours at 37°C. Leaves were subsequently destained for 48 hours in 50% ethanol, changing buffer once after 24 hours. Cleared leaves were mounted with glycerol on microscope slides and imaged.

### Epidermal cell size measurements and stomatal index

Images of uninfected adaxial leaf surfaces were taken at least 200 μm away from the midrib and the edges of the leaf, where at least 30 pavement epidermal cells could be counted. Area of cells was measured by tracing cell walls using the freehand shape tool in FIJI (https://imagej.net/Fiji) and using the measurement function to collect area of each shape. Stomatal counts were performed on the same images as for epidermal pavement cell size. Stomatal index was calculated as (number of stomata)/(total number of epidermal cells)*100 (as in Royer, 2001). Each leaf was imaged 3 times for counts and 3 leaves from different plants were used as biological replicates.

### Expression analyses

Tissue was immediately frozen in liquid nitrogen and stored at −80°C until extraction. RNA was extracted using Spectrum (Sigma-Aldrich) Plant Total RNA Kit from mock infected and infected tissues at 5 dpi according to manufacturer’s protocol. Residual genomic DNA was digested with DNase I (DNaseI, Qiagen). Purity and concentration of RNA was confirmed using the Nanodrop-1000 spectrophotometer (ThermoFisher Scientific). Complementary DNA (cDNA) was synthesized from 1μg RNA using High-Capacity cDNA Reverse Transcription Kit (ThermoFisher Scientific). cDNA was diluted 10x before use. Quantitative real-time PCR (qPCR) experiments were performed in a BioRad CFX96 (BioRad) using the iTaq Universal SYBR Green Supermix (Bio-Rad, USA), following kit instructions. Reactions contained 2μM primers with 2 μL of cDNA in 10 μL reactions. A negative control without cDNA was included in each set of reactions. For all genes, thermal cycling started with a 95°C denaturation step for 10 min followed by 40 cycles of denaturation at 95°C for 15 s and annealing at 56°C 30 s. Each run was finished with melt curve analysis to confirm specificity of amplicon. Three technical replicates were performed for each experimental set. Gene expression (fold change) was calculated using ACTIN2 as reference gene, and calculated using the Do My qPCR Calculations webtool (http://umrh-bioinfo.clermont.inrae.fr/do_my_qPCRcalc/; Tournayre et al. 2019). Each biological replicate consisted of a pool of leaves 7-9 from three plants. Statistically significant differences (p≤ 0.05; ANOVA and Tukey post-hoc test) are denoted by different letters for ANOVA, performed on each marker gene set. Primer sequences are provided in Supplemental Workbook S3.

### Lipid analyses

Tissue harvest and lipid extraction: Leaves 7-9 were harvested from mock infected and infected plants, washed of spores for collection, frozen in liquid nitrogen, and stored at −80°C until extraction. Extraction on all tissues was performed following lipase inactivation in 75°C isopropanol for 15 minutes according to Devaiah et al. (2006).

#### Electrospray ionization tandem mass spectrometry

The samples were dissolved in 1 ml chloroform. An aliquot of 10 to 20 μl of extract in chloroform was used. Precise amounts of internal standards, obtained and quantified as previously described (Welti et al. 2002), were added in the following quantities (with some small variation in amounts in different batches of internal standards): 0.36 nmol di14:0-PG, 0.36 nmol di24:1-PG, 0.36 nmol 14:0-lysoPG, 0.36 nmol 18:0-lysoPG, 2.01 nmol 16:0-18:0-MGDG, 0.39 nmol di18:0-MGDG, 0.49 nmol 16:0-18:0-DGDG, and 0.71 nmol di18:0-DGDG. The sample and internal standard mixture was combined with solvents, such that the ratio of chloroform/methanol/300 mM ammonium acetate in water was 300/665/35, and the final volume was 1.4 ml. Unfractionated lipid extracts were introduced by continuous infusion into the ESI source on a triple quadrupole MS/MS (API 4000, Applied Biosystems, Foster City, CA). Samples were introduced using an autosampler (LC Mini PAL, CTC Analytics AG, Zwingen, Switzerland) fitted with the required injection loop for the acquisition time and presented to the ESI needle at 30 μl/min. Sequential precursor and neutral loss scans of the extracts produce a series of spectra with each spectrum revealing a set of lipid species containing a common head group fragment. Lipid species were detected with the following scans: PG, [M + NH_4_]+ in positive ion mode with NL 189.0 for PG; lysoPG, [M – H]-in negative mode with Pre 152.9; MGDG, [M + NH_4_]+ in positive ion mode with NL179.1; and DGDG, [M + NH_4_]+ in positive ion mode with NL 341.1. The scan speed was 50 or 100 u per sec. The collision gas pressure was set at 2 (arbitrary units). The collision energies, with nitrogen in the collision cell, were +20 V for PG, +21 V for MGDG, +24 V for DGDG, and –57 V for lysoPG. Declustering potentials were +100 V for PG, +90 V for MGDG and DGDG, and –100 V for lysoPG. Entrance potentials were +14 V for PG, +10 V for MGDG and DGDG, and –10 V for lysoPG. Exit potentials were +14 V PG, +23 V for MGDG and DGDG, and –14 V for lysoPG. The mass analyzers were adjusted to a resolution of 0.7 u full width at half height. For each spectrum, 9 to 150 continuum scans were averaged in multiple channel analyzer (MCA) mode. The source temperature (heated nebulizer) was 100 C, the interface heater was on, +5.5 kV or −4.5 kV were applied to the electrospray capillary, the curtain gas was set at 20 (arbitrary units), and the two ion source gases were set at 45 (arbitrary units). The background of each spectrum was subtracted, the data were smoothed, and peak areas integrated using a custom script and Applied Biosystems Analyst software. The lipids in each class were quantified in comparison to the two internal standards of that class. The first and typically every 11th set of mass spectra were acquired on the internal standard mixture only. Peaks corresponding to the target lipids in these spectra were identified and molar amounts calculated in comparison to the internal standards on the same lipid class. To correct for chemical or instrumental noise in the samples, the molar amount of each lipid metabolite detected in the “internal standards only” spectra was subtracted from the molar amount of each metabolite calculated in each set of sample spectra. The data from each “internal standards only” set of spectra was used to correct the data from the following 10 samples. Finally, the data were corrected for the fraction of the sample analyzed and normalized to the sample “dry weights” to produce data in the units nmol/mgDW. Note that only species data with coefficient of variance (CoV) < 0.3 were included with CoV determined by independent measurements of identical samples. Statistically significant differences between genotype and treatment are denoted by different letters (tested by ANOVA and Tukey post hoc tests,* p ≤ 0.05).

#### Fatty acid methyl ester analysis

For FAME analysis, total lipid extracts were spiked with 25nmol pentadecanoic (C15:0) acid as internal standard. The sample was evaporated under a stream of nitrogen. Samples were resuspended in 1mL of 3M methanolic hydrochloric acid and heated at 78°C for 30 minutes. 2mL H_2_0 and 2mL hexane were added. Three hexane extractions were performed and dried down under a stream of nitrogen. Sample was then redissolved in 100μL hexane and analyzed on GC-FID (Agilent 6890N) after separating sample using DB-23 capillary column (column length - 60 m, internal diameter - 250 μm, film thickness - 0.25 μm). Carrier was helium gas at a flow rate of 1.5mL/min. The back inlet was operating at a pressure of 36.01 psi and 250 °C temperature. The GC oven temperature ramp was operated as follows: initial temperature of 150 °C hold 1min, increase at 25 °C/min to 175 °C. Then increase at 4°C/min to 230°C, hold 8 min. Total run time was 23.75 min. The FID was operated at 260 °C. The hydrogen flow to the detector was 30 mL/min, air flow was 400 mL/min and sampling rate of the FID was 20 Hz. Data processed using Agilent Chemstation software. Statistically significant differences between genotype and treatment are denoted by different letters (tested by ANOVA and Tukey post hoc tests,* p ≤ 0.05).

### Ploidy distribution of mesophyll cells at the powdery mildew infection site

Arabidopsis Col-0 and *acbp4-1* plants were infected with powdery mildew as above. At 5 dpi, 3 leaves from each plant were placed in 95% EtOH. Infected and mock uninfected leaves were vacuum infiltrated with 4’,6-diamidino-2-phenylindole (DAPI) (1 μg/mL) for twenty minutes at 200 Torr. Leaves were subsequently washed three times with PBS and stained with 1μg/ml Syto BC (Thermo Fisher Scientific) directly before visualization of leaves. DAPI quantification of mesophyll cell nuclei was achieved with the Zeiss LSM 710 confocal microscope. 50 slice Z-stacks were taken. Imaris Software v. 10.2 was used for 3D reconstruction to obtain DAPI fluorescent intensity maximums of the mesophyll cells in uninfected leaves and those underlying the haustorium. DAPI fluorescence values of epidermal pavement and mesophyll cell nuclei were normalized to that of nuclei of reference diploid guard cells from the same image after subtraction of background from each measurement as described in Chandran et al., 2013. Two independent experiments gave similar results, with ≥30 nuclei quantified for each time point and genotype. Significance of ploidy distribution with infection was assessed using nonparametric analysis of variance Kruskal-Wallis test (p≤0.05). Ploidy index of uninfected and infected *acbp4* plants calculated by Ploidy index = (%4C nuclei × 1) + (%8C nuclei × 2) + (%16C nuclei × 3) + (%32C nuclei × 4) + (%64C nuclei × 5); statistical differences were calculated by ANOVA and Tukey post-hoc test (p≤0.05).

### Hypocotyl Elongation in the Dark

Arabidopsis Col-0 and *acbp4-1* seeds bulked in parallel were surface-sterilized with 20% (v/v) bleach and stratified in 0.1% agarose (w/v) in the dark at 4°C for 2 days. Seeds were then plated on petri dishes containing 1/2 Murashige and Skoog pH 5.7, 0.5 g/L 2-(N-morpholino)ethanesulfonic acid (MES), 7 g/L agar, and 1 mL/L DMSO, exposed to light for 30 minutes, and then grown in the dark at 22°C for 4 days.

#### Hypocotyl length

measured from the base of the cotyledon to the root-shoot junction using the segmented line tool in ImageJ/FIJI. Significance tested by two-tailed t-test followed by Tukey’s HSD post-hoc test (*p ≤ 0.05, **p ≤0.01 ***p ≤0.001).

#### Cell Length

Epidermal cells of 4-day-old dark-grown Col-0 and *acbp4-1* seedlings were imaged using a 10x objective on a Leica AS LMD microscope. Three regions were imaged for each seedling: near the base (10% of the hypocotyl length above the root-shoot junction), at the midpoint, and near the apex (10% of the hypocotyl length below the cotyledons). Epidermal cell length was measured along the longitudinal axis of each cell, from the midpoint of the basal cell boundary to the midpoint of the apical cell boundary, using the segmented line tool in ImageJ/FIJI. Significance was tested by two-tailed t-test (*p ≤ 0.05, **p ≤ 0.01 ***p ≤ 0.001).

#### Ploidy Analyses

4-day-old dark-grown Col-0 and *acbp4-1* seedlings were stained using 1 μg/mL DAPI to visualize nuclei. Confocal Z-stack images were taken using a 20x objective on a LSM 710 confocal microscope at 3 regions for each seedling: near the base (10% of the hypocotyl length above the root-shoot junction), at the midpoint, and near the apex (10% of the hypocotyl length below the cotyledons). Nuclei were segmented in three dimensions using the Surfaces module in Imaris v9.5.0. Surfaces corresponding to non-epidermal nuclei in internal tissue layers were manually excluded. Total DAPI fluorescence intensity (intensity sum) and nuclear volume were extracted for each epidermal nucleus and log_2_-transformed before analysis.

Nuclear ploidy was inferred using an approach adapted from Russell et al. (2025). Log_2_-transformed intensity sum and nuclear volume were analyzed using a two-dimensional Gaussian mixture model with the mclust package (Scrucca et al. 2016) in R v4.4.2. A single model was fitted to nuclei pooled across genotypes and hypocotyl regions. Four clusters were specified based on the 2C, 4C, 8C, and 16C ploidy classes previously described in dark-grown Col-0 hypocotyls (Gendreau et al. 1998). Model fit was compared using Bayesian Information Criterion, and the best-supported model structure (VII) was used. The cluster with the lowest intensity sum and nuclear volume was designated 2C, with successively higher clusters designated 4C, 8C, and 16C. Significance in ploidy distribution between genotypes was tested by Pearson’s Chi-square test of independence (df=3; p ≤ 0.05).

## Supporting information

Supplementary Workbook S1

Supplementary Workbook S2

Supplementary Workbook S3

## ACKNOWLEDGEMENTS AND CONTRIBUTIONS

This work was supported by National Science Foundation (NSF) MCB-1617020 and USDA-National Institute of Food and Agriculture (NIFA) Hatch projects PLB-0223-H and CA-B-PLB-0330-H to MCW. CGT was supported in part by National Institute of Health (NIH) Genetic Dissection of Cells and Organisms Training Grant 5T32GM132022-04 & −05. JJ was supported in part by a UC Berkeley Arnon Fellowship. The lipid and FAME analyses described in this work were performed at the Kansas Lipidomics Research Center Analytical Laboratory. Instrument acquisition and lipidomics method development were supported by the NSF (including support from the Major Research Instrumentation program; most recent award DBI-1726527), K-IDeA Networks of Biomedical Research Excellence (INBRE) of NIH (P20GM103418), USDA-NIFA (Hatch/Multi-State project 1013013), and Kansas State University. Confocal microscopy and imaging analysis was performed at the RCNR Biological Imaging Facility at UC, Berkeley, supported in part by the NIH S10 program award number 1S10RR026866-01. The content is solely the responsibility of the authors and does not necessarily represent the official views of the NSF, NIH, or USDA-NIFA. All authors designed research and analyzed results. JJ, CGT, and KUW performed experiments. MCW, JJ, and CGT wrote the manuscript. We thank Dian Liu (UC Berkeley, Department of Plant & Microbial Biology) for critical review of the manuscript.

## SUPPLEMENTAL MATERIALS

- Supplemental Workbook S1: Gene sets and gene ontology enrichment results, supports Figure 2 and Discussion
- Supplemental Workbook S2: Lipid profiling, supports Figure 5
- Supplemental Workbook S3: Primers used in this study
- Supplemental Figure S1: *acbp4* T-DNA insertion lines and complementation, supports Figure 3
- Supplemental Figure S2: Cell ploidy at the PM infection site, supports Figure 6
- Supplemental Figure S3: Hypocotyl elongation in the dark, supports Figure 8

## LOCUS IDENTIFIERS

AT5G53470 - ACBP1

AT4G27780 - ACBP2

AT4G24230 - ACBP3

AT3G05420 - ACBP4

AT5G27630 - ACBP5

AT1G31812 - ACBP6

AT3G09840 - CDC48A

AT4G26100 - CKL1

AT4G28880 - CKL3

AT4G28860 - CKL4

AT1G48300 - DGAT3

AT3G16770 - EBP

AT4G22910 - FZR2/CCS52A1

AT3G20780 - HYP6

AT2G23430 - ICK1

AT5G11510 - MYB3R4

AT1G32640 - MYC2

AT5G10480 - PAS2

AT5G44420 - PDF1.2

AT5G58600 - PMR5

AT3G54920 - PMR6

AT2G14610 - PR-1

AT3G04720 - PR-4

AT3G27310 - PUX1

AT2G01650 - PUX2

AT5G25610 - RD22

AT1G49040 - SCD1

AT5G04470 - SIAMESE

AT5G05760 - SYP31

AT1G72260 - THI2.1

AT5G24770 - VSP2

AT3G56400 - WRKY70

