## Supplementary Figures and Tables for "Arabidopsis Acyl-CoA Binding Protein 4, ACBP4, functions in developmentally programmed endoreduplication"

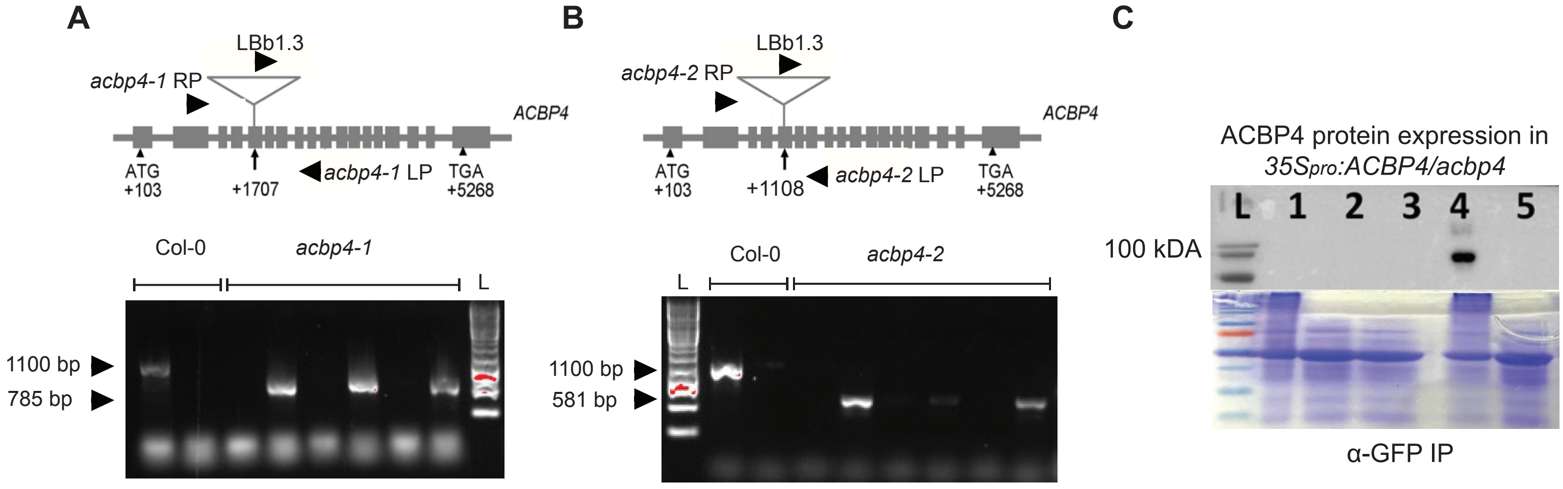


**Supplementary Figure S1:** ***acbp4*** **mutant line mapping and complementation.** (Supports Figure 3)

**A)** Gene map of *acbp4-1* allele (taken from Xiao et al. 2008), T-DNA insertion location, and genotyping primers. A representative genotyping gel for mutant plants with samples shown in adjacent lanes: the first lane utilizes primers that amplify the longer WT amplicon, while the second lane specifies the mutant, amplifying a shorter sequence from the T-DNA left border (LB) and the gene locus. Shown are representative WT Col-0 and *acbp4-1* plants screened for homozygosity. PCR product sizes are based on estimates from SALK T-DNA database (http://signal.salk.edu/tdnaprimers.2.html).

**B)** Gene map of *acbp4-2* allele,T-DNA insertion location, and genotyping primers. A representative genotyping gel for mutant plants with samples shown in adjacent lanes: the first lane utilizes primers that amplify the longer WT amplicon, while the second lane specifies the mutant, amplifying a shorter sequence from the T-DNA left border (LB) and the gene locus. L is DNA ladder. Shown are representative WT Col-0 and *acbp4-2* plants screened for homozygosity. PCR product sizes are based on estimates from SALK T-DNA database (http://signal.salk.edu/tdnaprimers.2.html).

**C)** Western blot showing ACBP4 expression in complemented $35S_{pro}$*:ACBP4/acbp4-1* line #2 (used in this manuscript). YFP-ACBP4 is visualized by Western using anti-GFP antibody (top) and loaded protein is visualized by Coomassie Stain of gel prepared in parallel (bottom).
Lane L is ladder, lane 1 is Col-0 protein extract enriched using GFP-antibody magnetic beads, lane 2 is crude Col-0 protein extract, lane 3 is crude protein extract from $35S_{pro}$*:ACBP4/acbp4-1* line #1, lane 4 is $35S_{pro}$*:ACBP4/acbp4-1* line #2 enriched using GFP-tagged magnetic beads, lane 5 is crude protein extract from $35S_{pro}$*:ACBP4/acbp4-1* line #2.

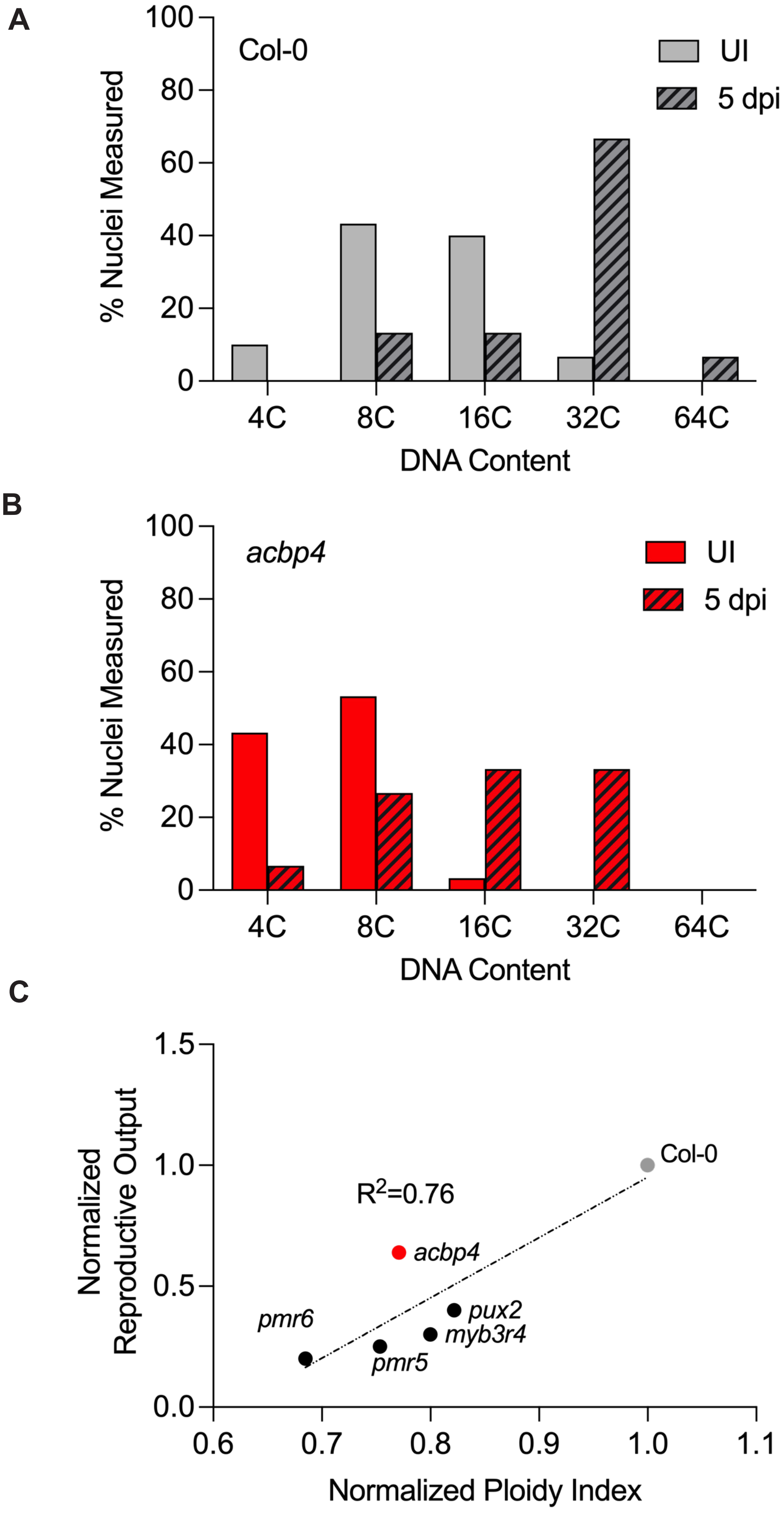

A
Col-0
UI
5 dpi
% Nuclei Measured
100
80
60
40
20
0
4C
8C
16C
32C
64C
DNA Content
B
acbp4
UI
5 dpi
% Nuclei Measured
100
80
60
40
20
0
4C
8C
16C
32C
64C
DNA Content
C
Normalized Reproductive Output
1.5
1.0
0.5
0.0
R2=0.76
Col-0
acbp4
pux2
myb3r4
pmr6
pmr5
0.6
0.7
0.8
0.9
1.0
1.1
Normalized Ploidy Index

**Supplementary Figure S2: Ploidy of mature leaf mesophyll cells is reduced in acbp4 compared to wild-type** (Supports Figure 6).
**A-B)** Ploidy distribution of the three mesophyll cells underlying the haustoria at 5 dpi compared to parallel cells from uninfected plant leaves for wild-type **(A)** and *acbp4* **(B)**. n ≥ 30 nuclei. One representative experiment is shown, the other experiment is shown in Figure 6. Both independent experiments yielded similar results. **C)** Normalized ploidy index plotted against normalized Gor reproductive output for *acbp4*, Col-0, and mutants tested in Chandran et al., 2012. $R^2 = 0.76$ determined by linear regression.

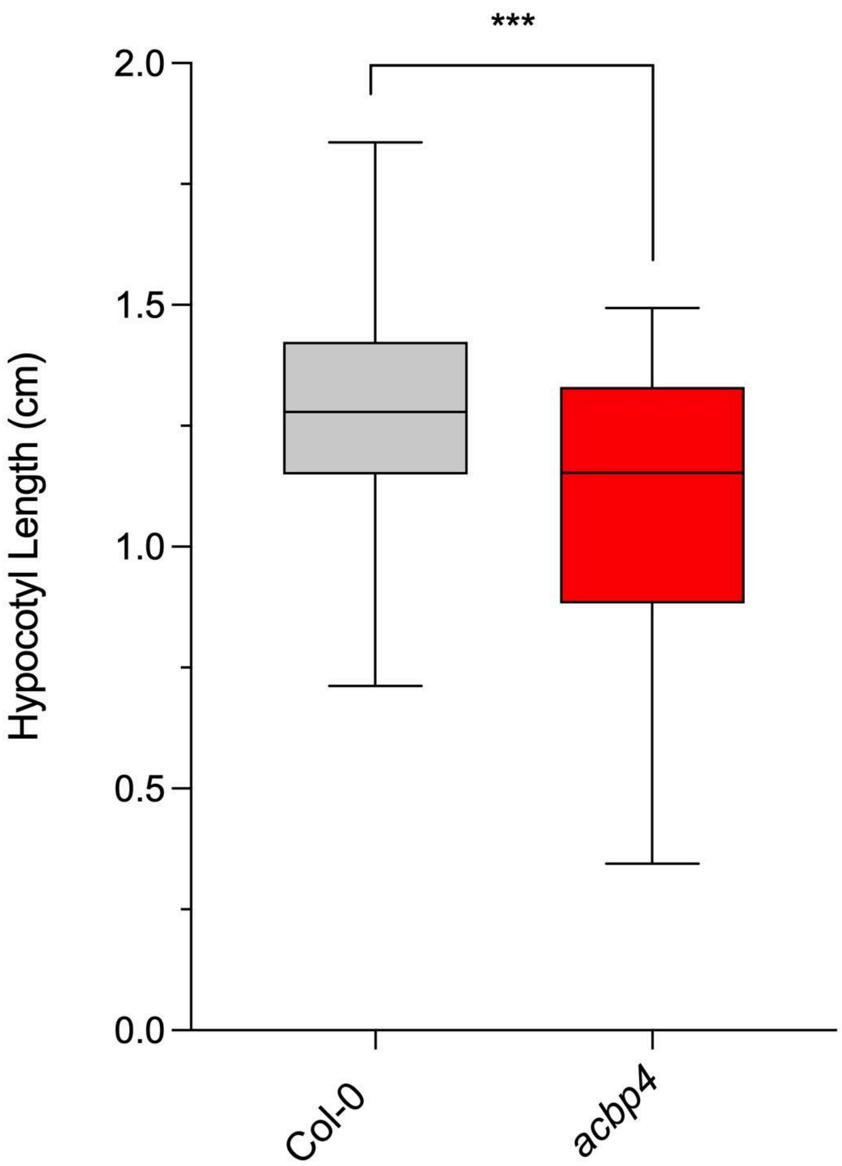


**Supplementary Figure S3: *acbp4* exhibits reduced growth in dark grown hypocotyls** (Supports Figure 8).

Hypocotyl lengths (cm) were measured for 4 day old Col-0 and *acbp4* seedlings grown on medium containing DMSO. In the box plots, the center line represents the median, the box limits indicate the upper and lower quartiles, and the whiskers extend to the highest and lowest values within 1.5 times the interquartile range. Sample sizes: Col-0 DMSO (n = 60), *acbp4*-DMSO (n = 57).One representative experiment is shown, the other experiment is shown in **Figure 8B**.
